# Climate risk for Italian habitats

**DOI:** 10.64898/2025.12.11.693144

**Authors:** Valerio Mezzanotte, Marta Cimatti, Sabina Burrascano, Moreno Di Marco

## Abstract

Climate change is a major driver of global biodiversity loss, and Europe is no exception with several areas exposed to high climate velocity and/or magnitude. Within the rapidly warming Europe, Italy is facing particularly high risk as part of the Mediterranean region with potentially dramatic consequences for its diverse habitat types. While species-level effects of climate risk are widely investigated, habitat-level exposure to climate change has rarely been assessed. This reduces the comprehensiveness of habitat state assessment under the Habitats Directive, and risks creating a blind spot on Protected Area effectiveness. Here we quantified the future (year 2050) climatic exposure of 139 EUNIS level-3 habitats across Italy, using two complementary metrics: analog velocity and multivariate magnitude. We found that median analog velocity was generally modest (median =0.15; Std = 0.41 km/y), and only a few habitats exceeded the critical velocity threshold of 0.5 km/yr. Instead, magnitude was typically high (median = 6.54; Std = 0.80) and 77% of habitat exceeded the critical threshold of 6.18. We also found a few habitats (2.15%) concurrently facing high velocity and high magnitude of climate change, mainly located in Apulia and Veneto. Mediterranean annual-rich dry grasslands and montane unvegetated inland shores were among the most exposed habitats (magnitudes ≈7.1–7.3; velocities ≈0.42–0.66 km/y), while cliff and mountain forest habitats in Sardinia and Sicily showed the lowest exposure. We urge countries to explicitly incorporate habitat-level exposure as part of national protection and restoration plans.

## Introduction

Climate change is a major threat to biodiversity, particularly in Europe, where warming was substantially faster than the global average in recent decades (Bilgili & Tokmakci, 2025; van Oldenborgh et al., 2009). Climate change impacts biodiversity at all levels (Scheffers et al., 2016), from genes to ecosystems to biomes, often by altering dramatically the distribution and composition of natural habitats. Much of the European conservation action is based on the Habitats Directive which focuses on natural habitats as “areas distinguished by geographic, abiotic and biotic features, whether entirely natural or semi-natural” and established the Natura2000 a network of protected areas. Nevertheless, while several studies on the impacts of climate change have been performed at the levels of biomes and species, (Allen et al., 2024; Bonannella et al., 2023; Loarie et al., 2009), only a handful have focused on habitats as described above (Heikkinen et al., 2022; Mücher et al., 2009). In this context, the need to investigate the climatic exposure of habitats is now more pressing than ever.

Statistical models are commonly used to assess species’ risk from climate change by linking their distributions to climatic variables. However reliable distributional data are lacking for the majority of species worldwide, making it impossible to fit reliable models (Garcia et al., 2014). The situation is more complicated for habitats, as habitat assessments often rely on expert judgment rather than systematic field data, resulting in an uncertain evaluation of their actual condition (AEE., 2019). The precise extent of many habitats remains poorly known, particularly outside the Natura 2000 network, hampering the implementation of comprehensive conservation strategies (Si-Moussi et al., 2025). Only recently Europe’s diverse range of habitats have been mapped at unprecedented resolution (Si-Moussi et al., 2025), providing new opportunities for spatially explicit assessments. It is also important to note that EUNIS habitat classes do not necessarily correspond uniquely to those used in the EU Habitats Directive. While these frameworks are related and sometimes directly overlap, each has been developed for distinct purposes and therefore applies different criteria and typological boundaries (John Rodwell et al., 2013).

Several studies have investigated climatic exposure of species at the European scale (Cimatti et al., 2025; Gillingham et al., 2024; Heikkinen et al., 2020; Lai et al., 2022; Nila & Hossain, 2019), but assessments explicitly targeting habitats remain scarce, especially for Italy. Despite Italy’s exceptional habitat diversity, also as part of the Mediterranean biodiversity hotspot, no systematic evaluation has yet quantified how different habitats across the country are exposed to future climate change. Existing national and regional studies have explored vegetation responses to climate change in specific areas (e.g. Gentilucci et al., 2023) or have addressed species (Maiorano et al., 2013) and communities associated with particular systems such as high-altitude mountain habitats (Ronchi & Brambilla, 2025). However, these analyses have narrow geographic or taxonomic scopes that do not provide a comprehensive and spatially explicit assessment across the full diversity of habitat types. Broader European assessments highlight the lack of integration of climate exposure metrics into habitat-level conservation planning (Cervellini et al., 2021),, as climate change may substantially alter the spatial patterns of habitat diversity. This makes a spatial analysis of habitats’ exposure to climate change critical for Italy, to inform effective conservation and adaptation strategies in one of Europe’s most biodiverse countries.

In this work we measure the climate exposure of Italian habitats (as defined by EUNIS). We provide spatially explicit assessments that allow us to discuss the most exposed habitats in terms of climatic velocity and magnitude, and the regional and national patterns of these exposures. These insights can inform the prioritization of management actions, including the expansion and adaptation of the protected area network and the plans for restoration, to ultimately strengthen biodiversity conservation efforts under accelerating climate change.

## Materials and Methods

### Input data

We focus our analysis in Italy, as it supports exceptionally high species and habitat diversity across multiple biogeographical regions (Alpine, Continental, Mediterranean). The Italian flora is one of the richest in Europe, with an extremely high share of endemic taxa (Bartolucci et al., 2018). This biotic diversity contributes to that of the Mediterranean biodiversity hotspot, and underpins a disproportionately large share of Europe’s conservation-priority taxa (Myers et al., 2000).

### EUNIS habitats

We mapped habitats in Italy using the European Nature Information System (EUNIS) habitat classification. EUNIS provides a comprehensive, hierarchical framework designed to cover the full spectrum of habitat types across Europe (Davies et al., 2004). EUNIS habitat classification is optimized for large-scale mapping and monitoring, including applications based on remote sensing that allow “wall-to-wall” spatial coverage (Si-Moussi et al., 2025). This functional focus distinguishes it from Potential Natural Vegetation (PNV) maps, which depict the hypothetical vegetation that would develop if the human impact ceased rather than the current distribution of habitats (Hengl et al., 2018; Somodi et al., 2021).

EUNIS habitat classification is a hierarchical system providing progressively more stringent habitat descriptions from Level 1 to Level 6. At the coarsest level (Level 1), major habitat formations such as wetlands, grasslands, and forests are identified. Narrower subdivisions are then introduced at each subsequent level. For the purposes of this study, the focus is on Level 3, which provides the most refined descriptions for terrestrial habitats in the revised classification from Chytrý and colleagues (Chytrý et al., 2020; Si-Moussi et al., 2025).

In our analysis we used data from Si-Moussi and colleagues (Si-Moussi et al., 2025) to define the presence of the most probable habitat class at 100 m resolution at EUNIS level 3 (Figure 1). Overall, the data includes 250 distinct habitat classes for Europe within nine broader formations (Table S1), and their predictions were evaluated against independent data from Austria, France (forests only) and the Netherlands (Si-Moussi et al., 2025). We excluded habitats that cover less than 1 square km across Italy, since their presence was too marginal for analysis. We also excluded cells covered by artificial surfaces or water (transitional waters, lakes, rivers), which accounted for 17,743 square km, and those cells without information for EUNIS level 3, for a total of 16,624 square km. This resulted in overall 34,367 square km excluded from the analysis, about 11% of the Italian surface, including 6% due to lack of information for EUNIS level 3.

**Figure 1.**
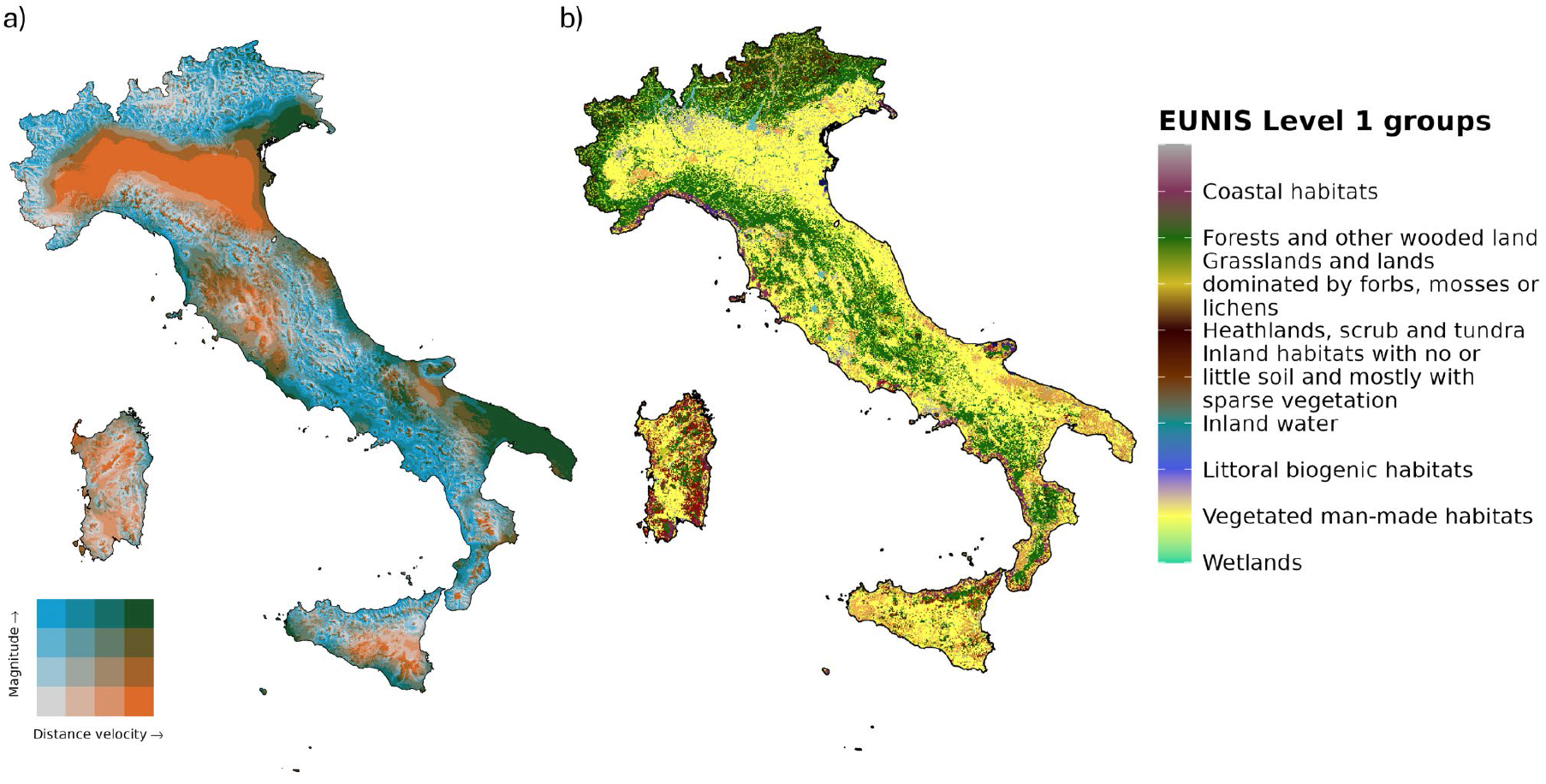
Climate risk for Italian habitats. a) Bivariate representation of climate metrics (analog/distance velocity and magnitude) in Italy under scenario SSP3-7.0 b) EUNIS habitats represented at level 2, color-coded according to their level 1 class.

### Climate exposure

We mapped climate exposure using two metrics of climatic change typically used to represent different dimensions of climate exposure for habitats and species (Brito-Morales et al., 2018; Cimatti et al., 2025; Garcia et al., 2014). We adapted two climate metrics calculated by Cimatti and colleagues (Cimatti et al., 2025) based on annual mean temperature and precipitation at 1 square km resolution, to quantify differences in climatic exposure of Italian habitats (Figure 1). We used the final model ensemble which results from the average of the three ESM analyzed for the two bioclimatic variables of interest (near surface mean annual temperature (2m) and total annual amount of precipitation). We focused on an intermediate Shared Socioeconomic Pattern - Representative Concentration Pathway (SSP-RCP) (O’Neill et al., 2016) SSP3-7.0. We used two complementary climate metrics: analog velocity (Brito-Morales et al., 2018; Hamann et al., 2015) and magnitude (Williams et al., 2007). Analog velocity evaluates the distance between regions that are projected to have similar climatic conditions in the future compared to the current climate in the focal location considered. This metric is useful to evaluate how much a species already on the edge of its climatic niche will need to move to track suitable climatic conditions, regardless of its actual possibility to do so (Brito-Morales et al., 2018). Magnitude quantifies the intensity of climate change at a given location, by measuring how much future climatic conditions depart from the historical baseline in multivariate climate space (Williams et al., 2007). This metric is adimensional and is scaled by baseline climate variability to summarize the intensity of multivariate climate change relative to past variability (in our case, for temperature and precipitation).

We refined data from Cimatti et al. (2025) by developing a new python-based algorithm (Mezzanotte & Mezzanotte, 2025) for calculating analog velocity in a more efficient way, which allowed us to extend the search buffer up to 5,000 km around each cell. As a result, we could minimize the number of no-analog cells (those with no climate analogue within the set search buffer), which we only found on the highest mountain tops (82 cells, equal to 0.026% of the Italian surface). Analog velocity elaboration is computationally demanding, as it may require scanning consistent parts of the study area to locate, for each focal cell, the nearest one with similar climatic characteristics. Our algorithm optimizes the search-path for analogs, going from the lowest Euclidean distance cells to the highest, and stopping once the first analog is found.

To aid interpretation of our results, we set biological threshold values for both metrics, to set levels of climate change which can be considered risky for biodiversity in a general sense. For climate velocity, we choose to consider 5 km per decade (0.5 km/y) used as a general dispersal proxy for plants by Warren and colleagues (Warren et al., 2013). For climate magnitude, we adopted the threshold of 6.18 set by Williams and colleagues (Williams et al., 2007) to identify novel climates in the mediterranean biome. We applied the same climate exposure values to all habitats found within the same cell (respectively 1km and 100 meters resolution).

## Results

### Climate exposure by habitat class

We found relatively low levels of climate velocity across Italian habitats, as only 2.7% out of 139 habitats reported median velocity values above the critical 0.5 km/y threshold. At the same time, we found generally high values of climate magnitude as 77% of habitats reported magnitude values above the 6.18 critical threshold (Figure 1, Table S1, Figures S2-S3).

Across the whole study area, habitats with the highest median exposures for both climate metrics were the *Mediterranean annual-rich dry grasslands* and the *Unvegetated or sparsely vegetated shore with mobile sediments in montane and alpine regions* (Table S1, Figure 2). These two habitats have an extension respectively of 45 and 2.3 square km (Table S1), and showed a velocity of 0.42 km/y and 0.66 km/y respectively and a magnitude of 7.3 and 7.1 (Table S1). Among widespread habitats (those covering an extent of at least 500 square km) the *Mediterranean lowland to submontane Pinus forest* and the *Mediterranean coniferous coastal dune forest* had the highest exposure, especially in terms of magnitude (6.31 and 7.04, respectively) (Table S1).

**Figure 2.**
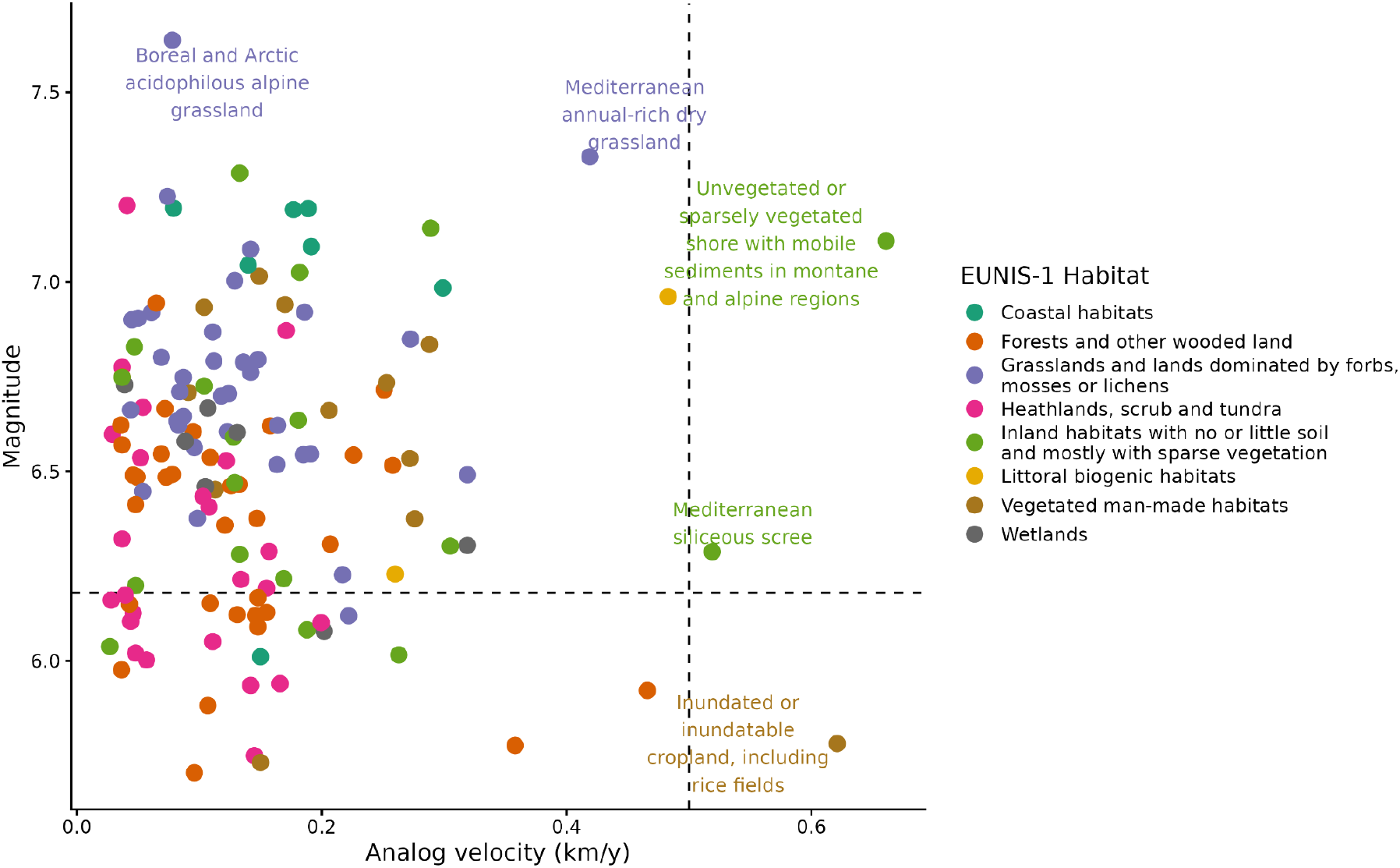
Median exposure level for analog velocity and magnitude for Italian habitats. Symbols represent individual EUNIS level-3 habitats, color-coded based on their EUNIS-1 level. The dashed lines show the thresholds of 0.5 km/y (vertical line) and of 6.18 (horizontal line).

The habitats exposed to high analog velocity and moderate magnitude are few, and relatively limited in extension. Among these are *Inundated or inundatable cropland, including rice fields* (covering about 2,800 km^2^, with a 0.62 km/y velocity and a 5.78 magnitude).

In terms of magnitude, the most exposed habitat were the *Boreal and Arctic acidophilous alpine grassland* (magnitude = 7.64) and the man-made habitat *Broadleaved fruit and nut tree orchards* (magnitude = 7.02). Among the most widespread habitats, a high median magnitude was found for *Temperate and submediterranean thermophilous deciduous forest* (covering about 30,000 km^2^, median magnitude of 6.54) and *Fagus forest on non-acid soils* (covering about 23,000 km^2^, median magnitude of 6.55).

Among the most exposed habitats at the national level there are those already identified at risk of collapse by the European Red List of Habitats (Table 1). Important examples include the *Temperate hardwood riparian forest*, and the *Mediterranean, Macaronesian and Black Sea shifting coastal dune*, which are respectively classified as Endangered and Vulnerable and are predicted to face high climate magnitude.

**Table 1.** European Red List of Habitats categories for the most exposed habitat across Italy, with their relative extension, median exposure values and upper relative EUNIS levels.

| EUNIS 1 | EUNIS 2 | EUNIS 3 | Median analog velocity | Median magnitude | RLE category EU28 | RLE category EU28+ | Area km<sup>2</sup> |
| --- | --- | --- | --- | --- | --- | --- | --- |
| Coastal habitats | Coastal dunes and sandy shores | Mediterranean, Macaronesian and Black Sea shifting coastal dune | 0.19 | 7.19 | Vulnerable | Vulnerable | 37.16 |
| Coastal habitats | Coastal dunes and sandy shores | Mediterranean and Black Sea sand beach | 0.18 | 7.19 | Near Threatened | Near Threatened | 52.34 |
| Forests and other wooded land | Broadleaved deciduous forests | Temperate hardwood riparian forest | 0.23 | 6.54 | Endangered | Endangered | 169.69 |
| Forests and other wooded land | Broadleaved deciduous forests | Carpinus and Quercus mesic deciduous forest | 0.16 | 6.62 | Near Threatened | Near Threatened | 1482.71 |
| Grasslands and lands dominated by forbs, mosses or lichens | Dry grasslands | Mediterranean annual-rich dry grassland | 0.42 | 7.33 | Near Threatened | Near Threatened | 45.13 |
| Grasslands and lands dominated by forbs, mosses or lichens | Dry grasslands | Continental dry grassland (true steppe) | 0.22 | 6.23 | Near Threatened | Near Threatened | 77.13 |
| Grasslands and lands dominated by forbs, mosses or lichens | Dry grasslands | Cryptogam- and annual-dominated vegetation on siliceous rock outcrops | 0.19 | 6.55 | Vulnerable | Vulnerable | 6.41 |
| Grasslands and lands dominated by forbs, mosses or lichens | Dry grasslands | Mediterranean to Atlantic open, dry, acid and neutral grassland | 0.18 | 6.54 | Vulnerable | Near Threatened | 68.74 |
| Grasslands and lands dominated by forbs, mosses or lichens | Dry grasslands | Oceanic to subcontinental inland sand grassland on dry acid and neutral soils | 0.16 | 6.62 | Endangered | Endangered | 19.54 |
| Heathlands, scrub and tundra | Maquis, arborescent matorral and thermo-Mediterranean scrub | Thermomediterranean arid scrub | 0.16 | 6.29 | Vulnerable | Vulnerable | 122.58 |

### Spatial variation of habitat risk across regions

The climatic exposure of habitats varied significantly across Italian regions (Figures S2-S22), with Apulia and Veneto showing the highest level of exposure for both climate metrics, Emilia-Romagna showing some of the highest values of velocity, and Campania some of the highest magnitudes. For example, the *Alpine and subalpine ericoid heath* habitat is more exposed to magnitude in Veneto than in Lombardy (6.99 vs. 6.04) despite having comparable spatial extension (245 and 161 square km) (Table S2). Several habitats show different exposure to either velocity or magnitude or both although their similar extents across regions: *Fagus forests on non-acid soils* from Emilia-Romagna are much more exposed to velocity than those in Lombardy (0.16 vs 0.04 km/y) but the opposite is true for magnitude (Table S2). The *Mediterranean maquis and arborescent matorral* from Apulia is more exposed than the same habitat in Calabria to both velocity (0.48 vs 0.06 km/y) and magnitude (7.43 vs 6.90). The *Temperate subalpine Larix, Pinus cembra and Pinus uncinata forest* from both Trentino-South Tyrol and Piedmont show low levels of velocity (0.05 for both of them), but different levels for magnitude (7.03 vs 6.07). The *Mediterranean lowland to submontane Pinus forest* from Apulia and Sardinia differ for both velocity (0.67 vs. 0.17 km/y) and magnitude values (7.09 vs. 5.79). The *thermomediterranean arid scrub* from Sicily and Sardinia show large variation in magnitude (6.80 vs. 5.73).

On the other hand, some habitats have high risk values across different regions, such as the *Mediterranean coniferous coastal dune forest* which had high magnitude (6.94-7.39) across all regions with highest coverage (Liguria, Calabria, Apulia, Tuscany; Figure 3). The same is true for *Temperate and submediterranean thermophilous deciduous forest*, even if this had slightly lower magnitude (6.05-6.67; Table S2). We also found regional risk for some of the habitats with a high conservation value. For example, the *Temperate subalpine Larix, Pinus cembra and Pinus uncinata forest* is classified as a priority habitat* in the Habitats Directive (codes 9420 and 9430) if found on calcareous or gypsum substrates, and we found that more than 75% of the habitat in Trentino-South Tyrol is above the 6.18 critical threshold for magnitude (Figure 3b).

**Figure 3.**
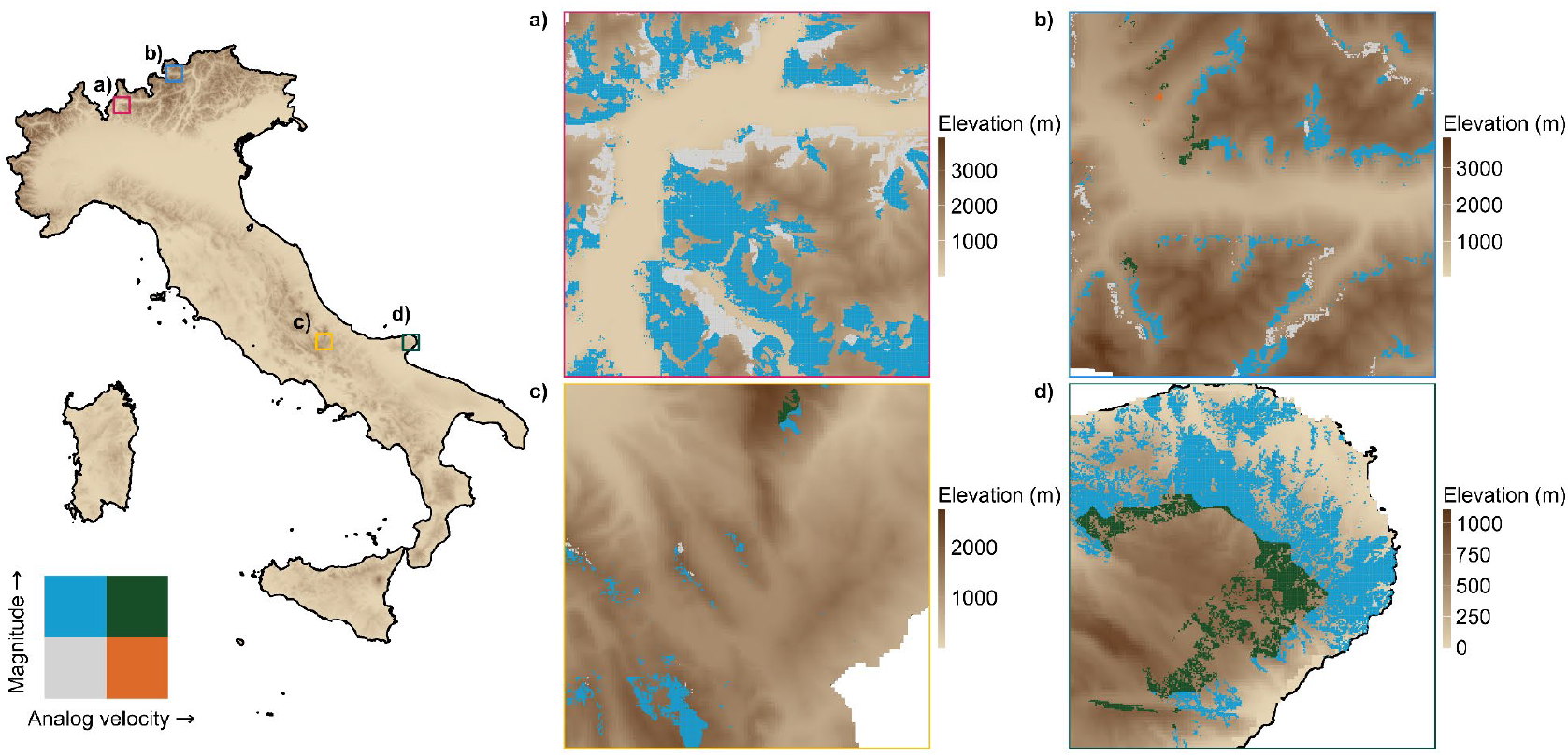
Exposure level for analog velocity and magnitude for EUNIS 3 level habitat, using 0.5 km/y and 6.18 as thresholds. Colors represent the presence of the habitat of interest and its climatic exposure. Blue represents areas exposed to magnitude values higher than 6.18, orange represents areas with velocity values higher than 0.5 km/y, green highlights areas where both thresholds are exceeded and the grey areas where both metrics are below these critical thresholds. We show the exposure of a variety of habitats: in a) *Fagus forest on non acid soil* in Lombardy, in b) *Temperate subalpine Larix, Pinus cembra and Pinus uncinata forest* in Trentino-South Tyrol, in c) *Temperate acidophilous alpine grassland in Abruzzo*, in d) *Mediterranean coniferous coastal dune forest in Apulia*.

The regional habitats with the highest velocity were the *Cyrno-Sardean oromediterranean siliceous dry grassland* of Sardinia, with a median value of 3.72 km/y (16.9 times higher than national average; Table S3), followed by the *Temperate acidophilous alpine grassland* of Calabria and Tuscany, respectively with 2.76 km/y (34.5 times higher than national average) and 1.96 km/y (24.5 times higher than national average).

For magnitude, the *Spartium junceum scrub*, the *Mediterranean upper-mid saltmarsh and saline and brackish reed, rush and sedge bed* and the *Alpine and subalpine calcareous grassland of the Balkans and Apennines*, all from Apulia, reported the highest values, all above 7.5 (Table S3). We show some examples of regional habitats with their relative local exposure levels (Figure 3). Overall, 12.1% of the regional habitats analyzed reported values above 0.5 km/y for analog velocity and 6.18 for magnitude (Table S3; Figures S15, S22).

## Discussion

Our analysis provides the first national-scale assessment of climatic exposure for Italian EUNIS habitats, using two complementary climate metrics. We found a high share of the study area is undergoing intense magnitude of climate change, yet with a relatively low velocity. These general outcomes, however, show a certain degree of variation across habitat types and regions, thus pointing to the need for both national and regional targets to be pursued locally in a coordinated way. This is true also because habitat environmental conditions other than climate may limit the possibility of the species dominating different habitats to find a favorable site. Many of the habitats facing high climate risk are linked to specific substrates or topographic conditions, e.g., siliceous bedrocks, mobile sediments, cliffs. These lithological and morphological heterogeneity is to be accounted for at more local scales than climate, as it was reported to drive landscape and biodiversity patterns at finer scales than climate since the earliest studies on ecological classifications (Klijn & de Haes, 1994).

We found that, while analog velocity was relatively limited across most habitats, magnitude diffusely exceeded the critical threshold identified in the literature, indicating that the great majority of habitats in Italy are projected to experience climatic conditions that deviate substantially from their historical baseline. This suggests that although the required spatial displacement to track climate analogs may be low to moderate for many habitats, the intensity of local changes will be considerable. Together, these findings highlight the importance of interpreting velocity and magnitude jointly, as they capture different but complementary facets of exposure.

A few habitats had high (above threshold) exposure to both climate magnitude and velocity, these require urgent assessment of their distribution and conservation state. Both *Annual Mediterranean grasslands* and *Inland shores with mobile sediments* are in this condition, and are extremely challenging to map. In fact, they occur in small and dynamic patches whose distribution may shift rapidly in response to ecological succession and disturbances (for the *Annual Mediterranean grasslands*), and to the modification of shores after sediment erosion and deposition (for the *Inland shores with mobile sediments)*.

Many other habitats, much more widespread, are exposed to high magnitude but low or moderate velocity. This is the case of the *Thermophilous deciduous forest*. Although the low velocity value obtained for this habitat may not lead to an urgent threat, the dominant species of these habitats – mostly deciduous oaks such as *Quercus cerris, Q. pubescens, Q. frainetto –* have the highest regeneration densities between 140 and 330 meters from the seed source (Axer et al., 2021), with a mean dispersal distance by birds of 115 meters (Kurek et al., 2019). These values are barely above the median value of analog velocity for this habitat (0.11 km/y), meaning that climate velocity might outpace seed dispersal. Moreover, oak species, like most other tree species, take many years to reach sexual maturity (ca. 30-50 years), which means an expansion of habitats will take at least decades to complete. An even worse scenario is the one for *Fagus forest on non-acid soils*, whose dominant species, the beech (*Fagus sylvatica*), was reported to have an average dispersal distances of just 12-25 meters (Bontemps et al., 2013), with a similar maturity age as oak species. Moreover, seed production in beech and oak forests follows multi-year cycles linked to climate, i.e., mast cycles. These cycles are renownedly altered by climate change, which was found to increase mast frequency at the expense of growth in beech (Hacket-Pain et al., 2025) with evidence of a concomitant increase of seed predation pressures (Bogdziewicz et al., 2020).

We found that several of the most exposed habitats are included in the European Red List of Habitats. One of these is the Temperate hardwood riparian forest (Table 1), a habitat that underwent a 80% loss of its European extent during the past 200 years (Naiman et al., 1993). The vulnerability of this habitat is linked to a high degree of fragmentation often induced by land conversion for agriculture or urbanization, and often occurring together with ditching, addition of pesticides and fertilizers (Riis et al., 2020). Fragmentation also worsens the impact of biological invasions by other woody and non-woody species, which is promoted by the direct connection to water systems, and by the disturbance-prone character of this habitat (Richardson et al., 2007). Climate change could act synergistically with these risks and further limit the habitat persistency in the long term: because of its prevalence in floodplains the habitat will be subject to intermediate values of velocity (0.23 km/y) and will also be subject to above the critical threshold value (6.54) for magnitude, potentially impairing its ability to persist in place.

Exposure patterns varied markedly among habitats and across regions, implying that the same habitat may be at risk in one region but safe in another one, underscoring the need for subnational conservation strategies. For example, every habitat found in Apulia has a higher climate magnitude compared to almost every other region. The *Cyrno-Sardean oromediterranean siliceous dry grassland* instead shows much higher velocity in Sardinia than elsewhere. This habitat is found in the slopes of high mountains, whose climate analogs are rare in the island of Sardinia.

Several habitats facing high climate risk according to our analysis are already classified as having an unfavorable conservation status, such as the Mediterranean lowland to submontane Pinus forest (Habitat 9540 in the Habitats Directive; ISPRA, 2021). This is due to their inadequate management of these forests, which are often planted and highly homogeneous (Biondi et al., 2010), i.e., high density of trees of the same species, and thus more vulnerable to disturbances that are frequent in this habitat and amplified by climate change, such as fire and pest disturbances.

The ecological interpretation of climate metrics depends strongly on species traits and ecological processes. Responses to both velocity and magnitude are highly species-specific and mediated by life-history traits, climatic niche breadth, demographic rates, landscape connectivity, and biotic interactions. When looking at animal species, several authors have compared climate velocity with the dispersal ability of species (Newbold, 2018; Warren et al., 2018). For plants and, consequently, habitats there are no broadly accepted thresholds. Empirical and paleoecological studies report very large variation in migration and dispersal rates of plants, ranging from only a few meters to several kilometers per year under certain modelled scenarios (McLachlan et al., 2005), with rare long-distance dispersal events that can sometimes drive faster realized range shifts, a phenomenon often referred to as Reid’s paradox (Clark et al., 1998). For instance, an analysis of 140 plant species found an average postglacial migration rate of 0.12 km/year, while average potential future migration rate was predicted as 1.6 km/year (Cunze et al., 2013). Interestingly, resource allocation in plant organs lead to proven trade-offs, e.g., a species cannot deploy both a large light-capturing area and strongly reinforced and long-lasting leaves, or large and nutrient rich seeds without producing fewer of them (Laughlin, 2023; Westoby, 1998). These tradeoffs imply that long-distance colonizers producing many light seeds are mostly species with short-lived, cheaply constructed leaves and rapid growth rates, adapted to exploit transient, resource-rich environments. Conversely, species investing in fewer, heavier seeds tend to be more competitive in stable, resource-poor habitats, often exhibiting slow growth, high tissue density, and long leaf lifespans. Thus the latter species, i.e., stress tolerant species, are those that we may expect to be more impacted by high velocity, and are also those typically developing in the mountain and alpine grasslands subjected to the highest velocity values. Plants’ response to climate magnitude depends on factors such as niche breadth, adaptive capacity, and dispersal ability. Williams et al. (2007) derived a conservative threshold of 6.18 specific to the Mediterranean biome to define novel climates (which we adopted in our study). Values exceeding these thresholds suggest that future climates will fall outside the range of historical variability and are unlikely to remain within the tolerance envelopes of species and communities adapted to past conditions. Under such scenarios, local persistence becomes increasingly uncertain unless populations can undergo rapid evolutionary or phenotypic adaptation, exhibit strong plasticity, or benefit from conservation interventions such as assisted migration.

The pervasive exceedance of magnitude thresholds across Italian habitats suggests that climate change will affect most habitats, with some facing particularly high risks. Climate change threatens virtually all habitats, challenging the Habitats Directive framework in the medium and long term. The first key implication of our study is that climate change will not hit all habitats in the same ways: spatial heterogeneity will be a crucial aspect, even within the same habitat. Clearly, this implies tailoring conservation measures to each specific habitat based on their different exposure, vulnerability and conservation objectives. This has direct implications for the Nature Restoration Law, knowing the exposure level of each habitat is a fundamental step to identify the habitat most in need of restoration actions to mitigate climate change effects. Second, we highlight that only three habitats in the European Red List of Habitats have been classified as threatened based on future habitat loss (criterion A2), despite dozens of them being known to be threatened by climate change. Moreover, many habitats that we identify as highly exposed to climate change are already classified as impacted by other threats, with high risk of synergistic effects among threats (Brook et al., 2008; Mantyka-Pringle et al., 2013).

Conservation and restoration strategies should recognize and explicitly account for climate change in the long term before implementing any kind of actions, knowing that refugia represent a safe investment for the future of biodiversity and that hotspots on the other hand may need heavier investment and interventions than other areas. We recommend prioritizing restoration in climatically stable areas wherever possible, and include climate adaptation actions in all cases where restricted habitats are expected to face high climate magnitude. Landscape planning should enhance habitat connectivity and adjust disturbance regimes towards habitat conservation and improved resilience. These actions often involve adjustments to fire and grazing, improved seed sourcing and restoration practices to increase heterogeneity and biodiversity both in grasslands (Bernath-Plaisted et al., 2025) and forests, in which promoting more close-to nature silvicultural approaches, such as structurally rich forest stands with diverse species composition contribute to increased resistance and resilience of forests to climate change (Nabuurs et al., 2018).

## Conflict of interest

The authors declare no conflicts of interest.

## Acknowledgements

*We acknowledge funding by The European Union–NextGenerationEU as part of the National Biodiversity Future Center, Italian National Recovery and Resilience Plan (NRRP) Mission 4 Component 2 Investment 1*.*4 (CUP: B83C22002950007)*.

## Notes

https://github.com/valerio3mezzanotte/analog_velocity_python/blob/main/README.md

### Competing Interest Statement

The authors have declared no competing interest.

