## Supplementary material for "Climate risk for Italian habitats": Supplemetantary_Material

### Supplementary results:

#### Analog velocity adaptation results

We showed that the 82 cells which found no climate analog within Cimatti et al., 2025 study', showed an analog within the 3000 km search radius (about 2700 km distance for farthest analogue).

##### Table S1. Median values for climatic exposure and area covered by each EUNIS level 3 habitat for Italy

Attached as a separate file

##### Table S2. Median differences in climatic exposure for habitats found in multiple regions across Italy.

Attached as a separate file

##### Table S3. Median values for climatic exposure and area covered by each EUNIS level 3 habitat for each Italian region

Attached as a separate file

**Climate exposure for Italy under Mediterranean threshold**

Analog velocity & magnitude under SSP3-7.0

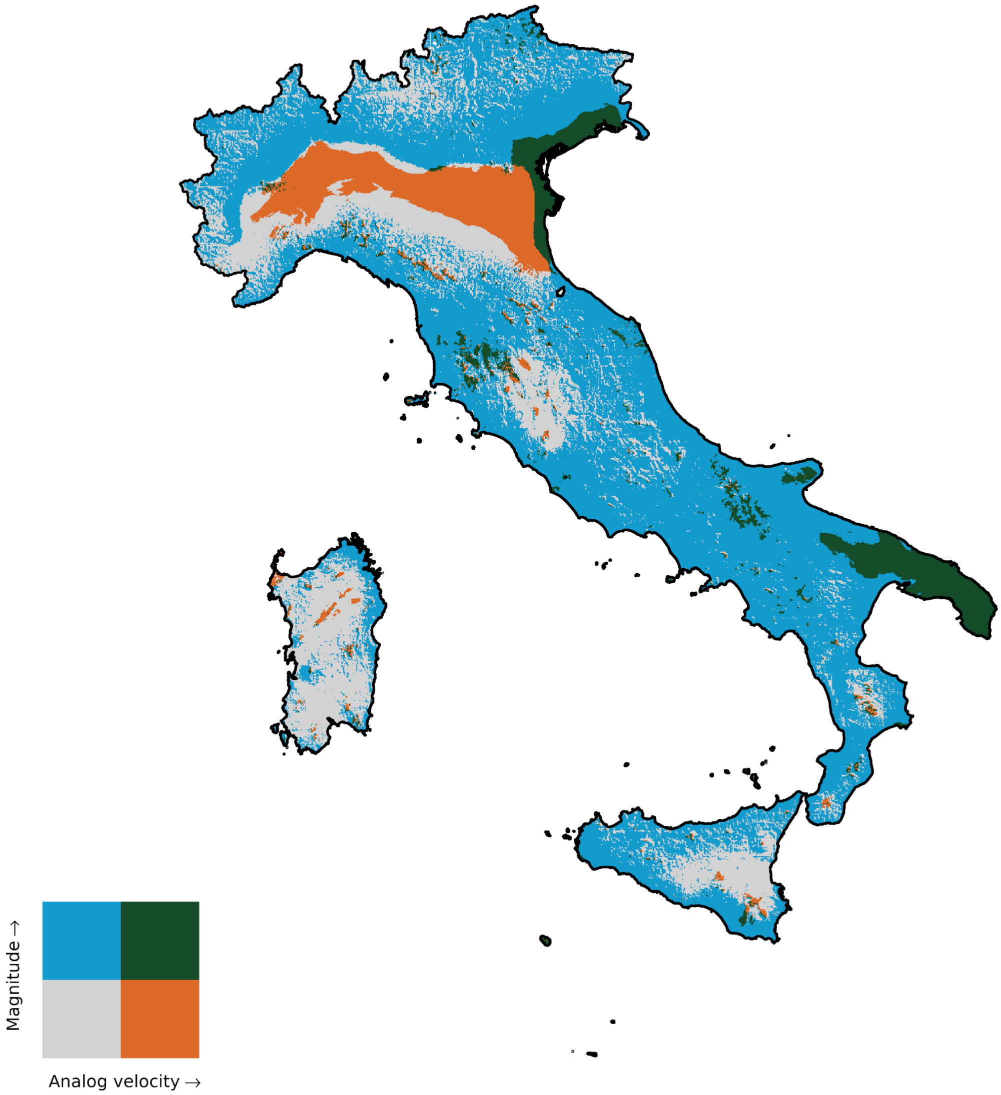

Figure S1. Climate exposure in Italy under SSP3-7.0, binarized to match significant biological thresholds. For analog velocity it is 0.5 km/y (equal to 5  km shift per decade) and for climate magnitude it is here 6.18.

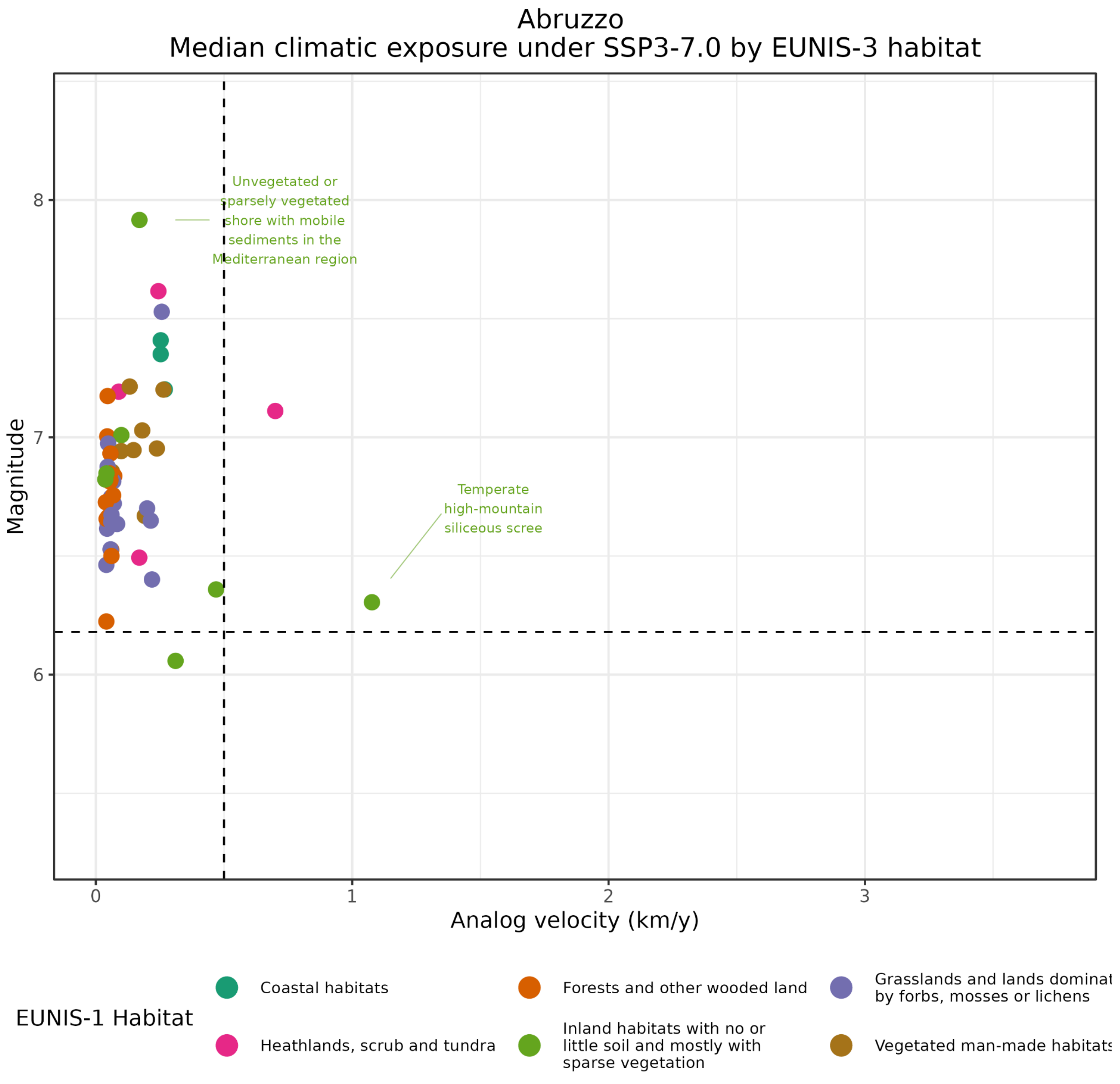


Figure S2. Median climate exposure for each EUNIS level 3 habitat for Abruzzo under SSP3-7.0

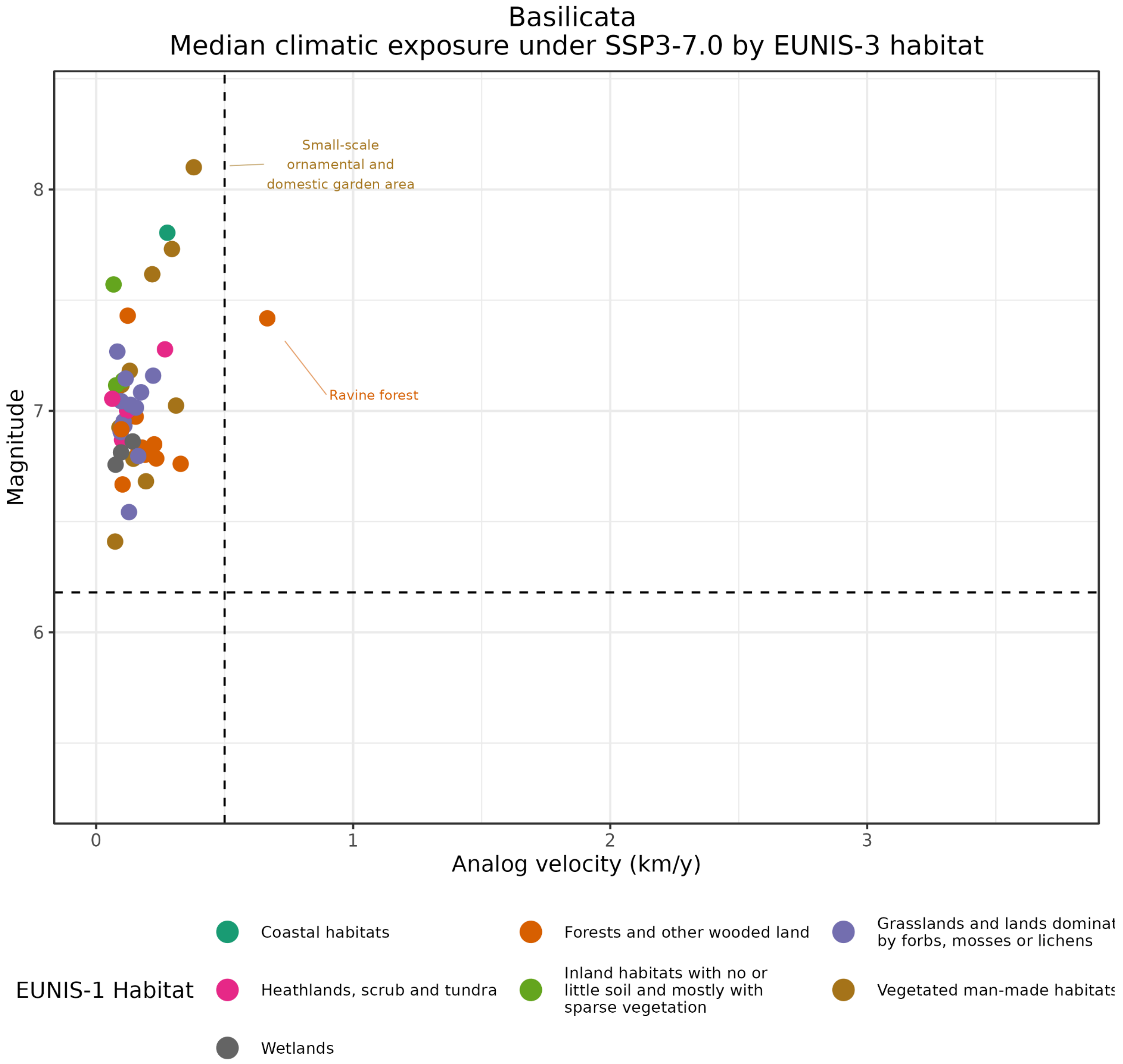
Basilicata
Median climatic exposure under SSP3-7.0 by EUNIS-3 habitat
Small-scale
ornamental and
domestic garden area
Ravine forest
8
7
6
Magnitude
0
1
2
3
Analog velocity (km/y)
EUNIS-1 Habitat
Coastal habitats
Forests and other wooded land
Grasslands and lands dominat
by forbs, mosses or lichens
Heathlands, scrub and tundra
Inland habitats with no or
little soil and mostly with
sparse vegetation
Vegetated man-made habitats
Wetlands

Figure S3. Median climate exposure for each EUNIS level 3 habitat for Basilicata under SSP3-7.0

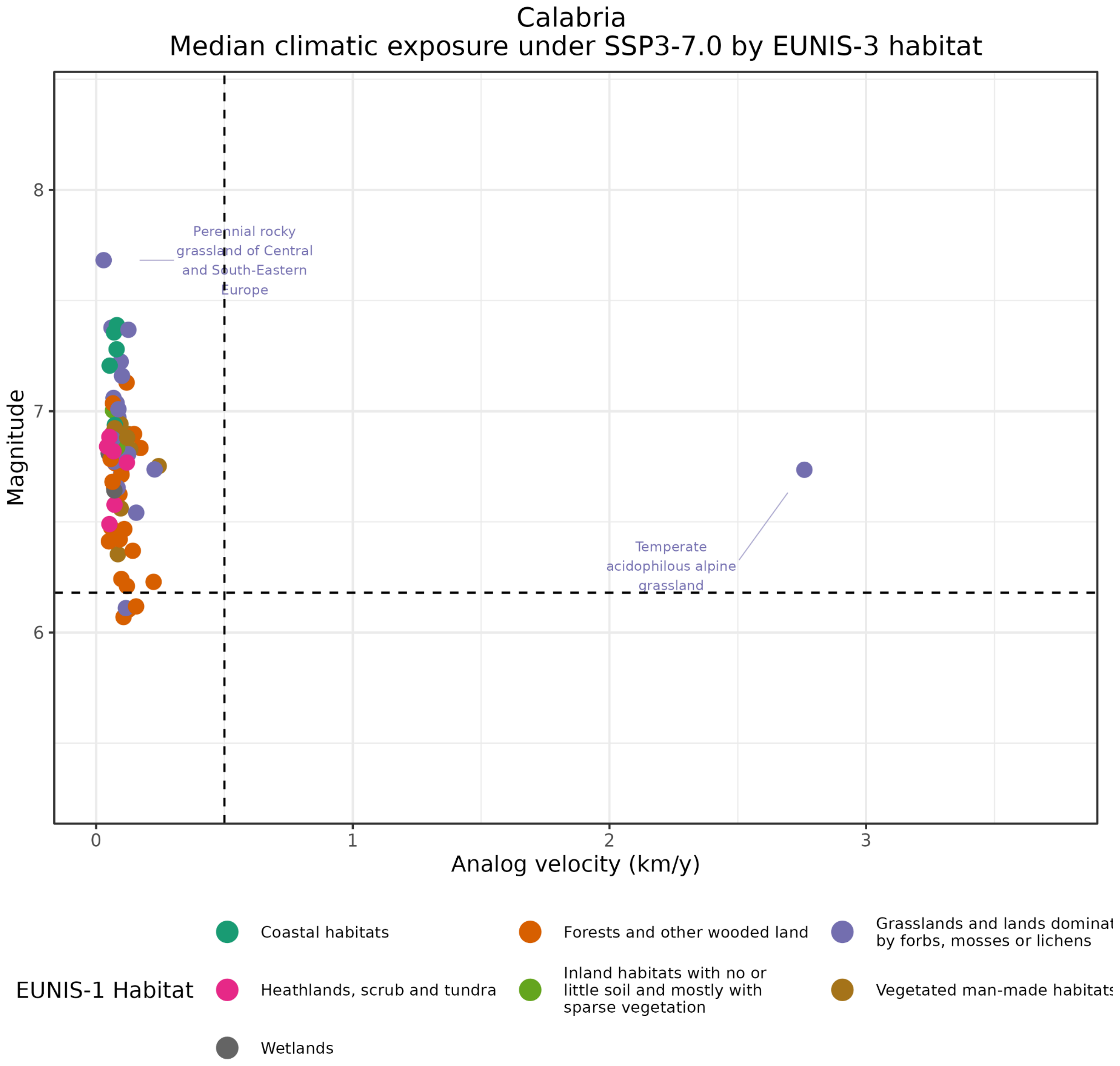

Figure S4. Median climate exposure for each EUNIS level 3 habitat for Calabria under SSP3-7.0

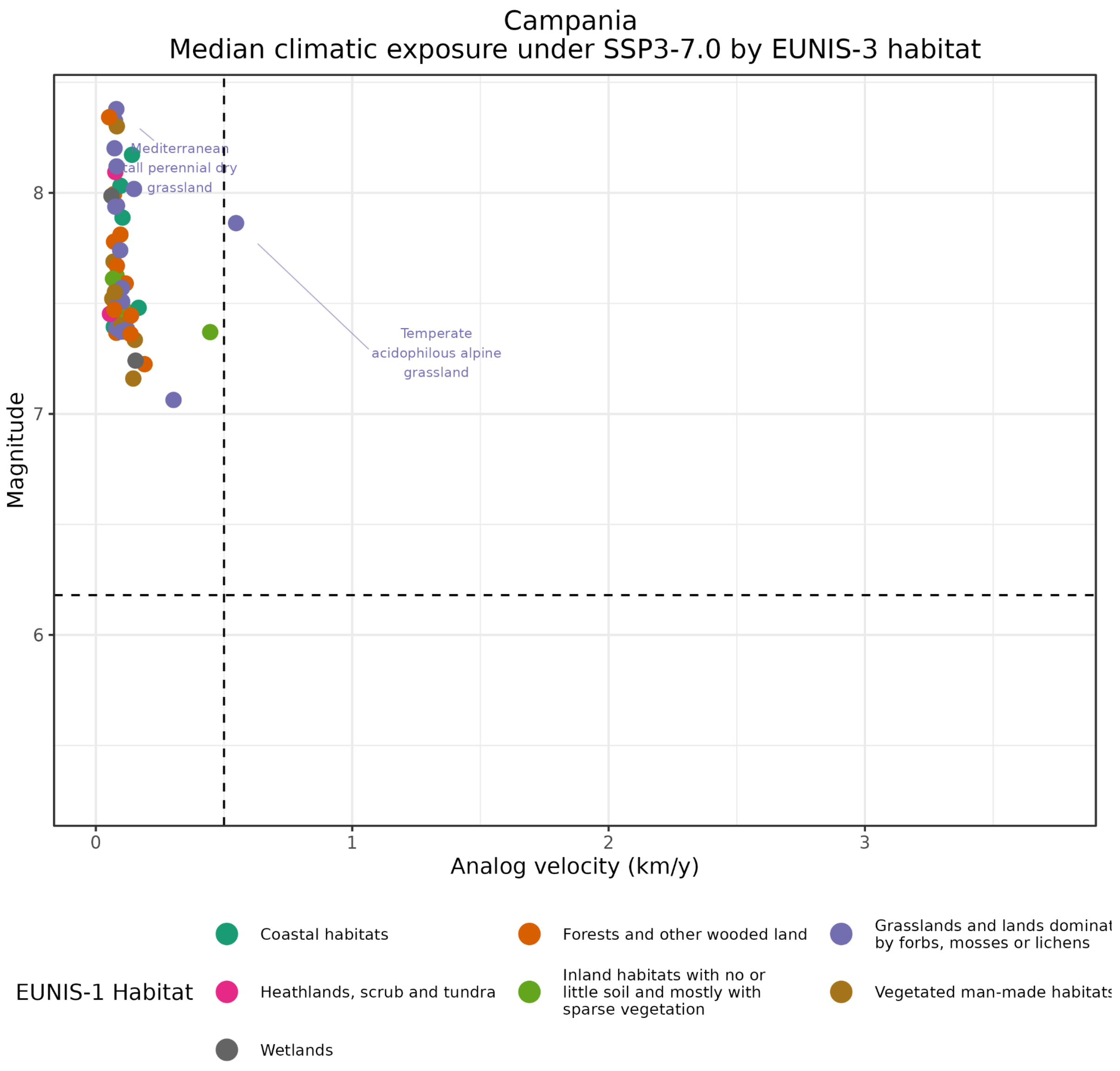


Figure S5. Median climate exposure for each EUNIS level 3 habitat for Campania under SSP3-7.0

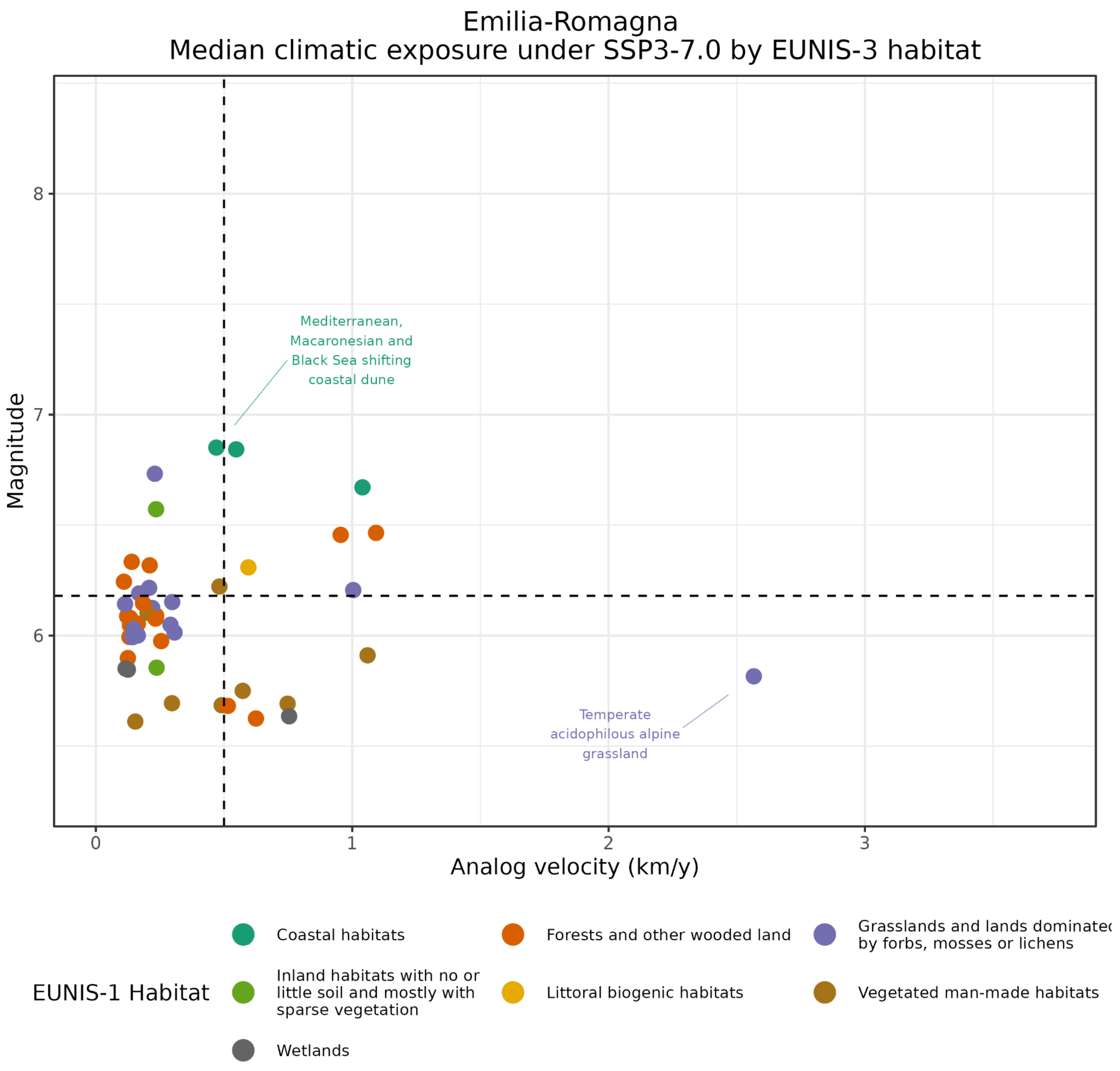
Emilia-Romagna
Median climatic exposure under SSP3-7.0 by EUNIS-3 habitat
Magnitude
8
7
6
Mediterranean, Macaronesian and Black Sea shifting coastal dune
Temperate acidophilous alpine grassland
0
1
2
3
Analog velocity (km/y)
EUNIS-1 Habitat
Coastal habitats
Forests and other wooded land
Grasslands and lands dominated by forbs, mosses or lichens
Inland habitats with no or little soil and mostly with sparse vegetation
Littoral biogenic habitats
Vegetated man-made habitats
Wetlands

Figure S6. Median climate exposure for each EUNIS level 3 habitat for Emilia-Romagna under SSP3-7.0

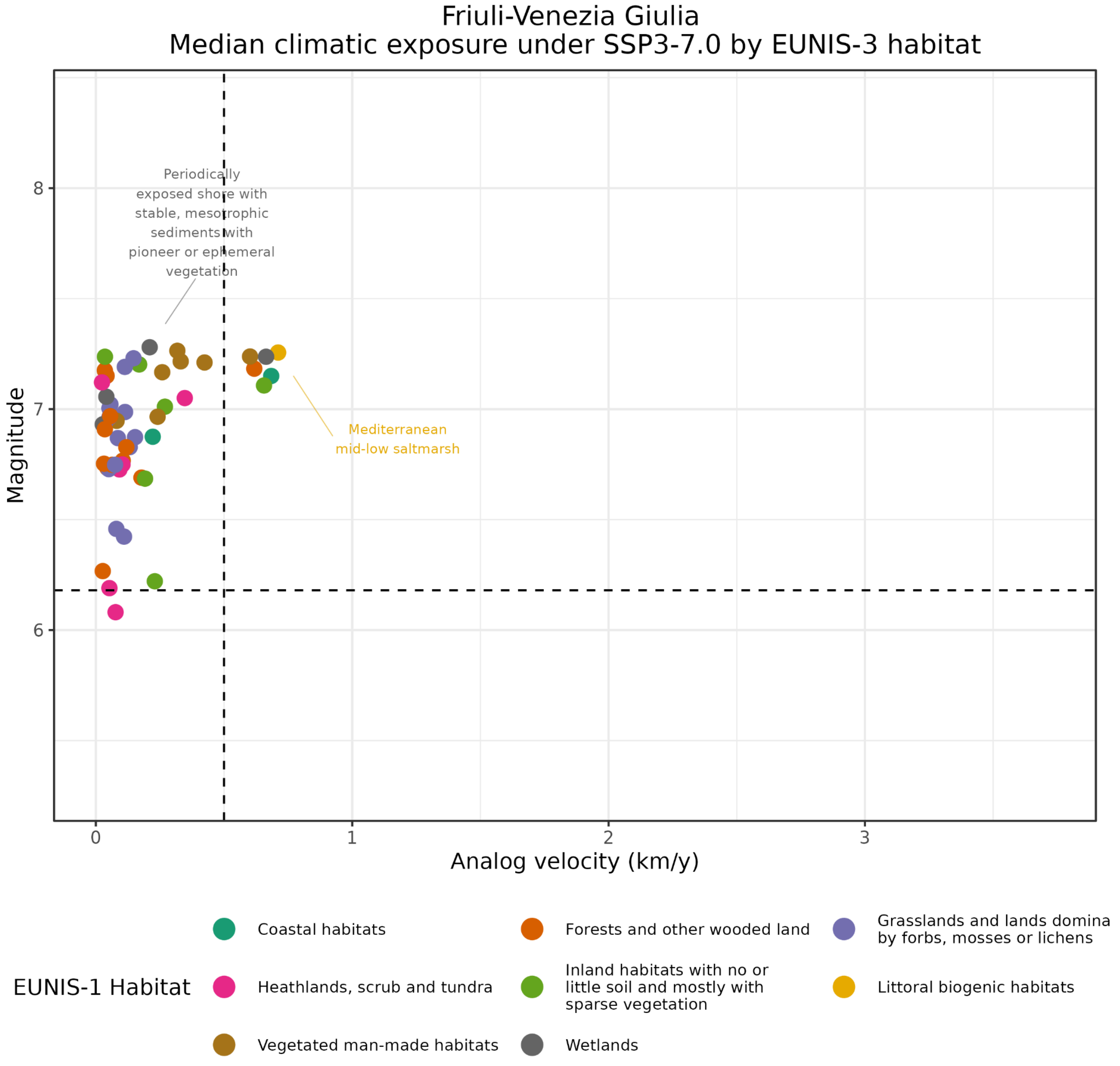

Figure S7. Median climate exposure for each EUNIS level 3 habitat for Friuli Venezia Giulia under SSP3-7.0

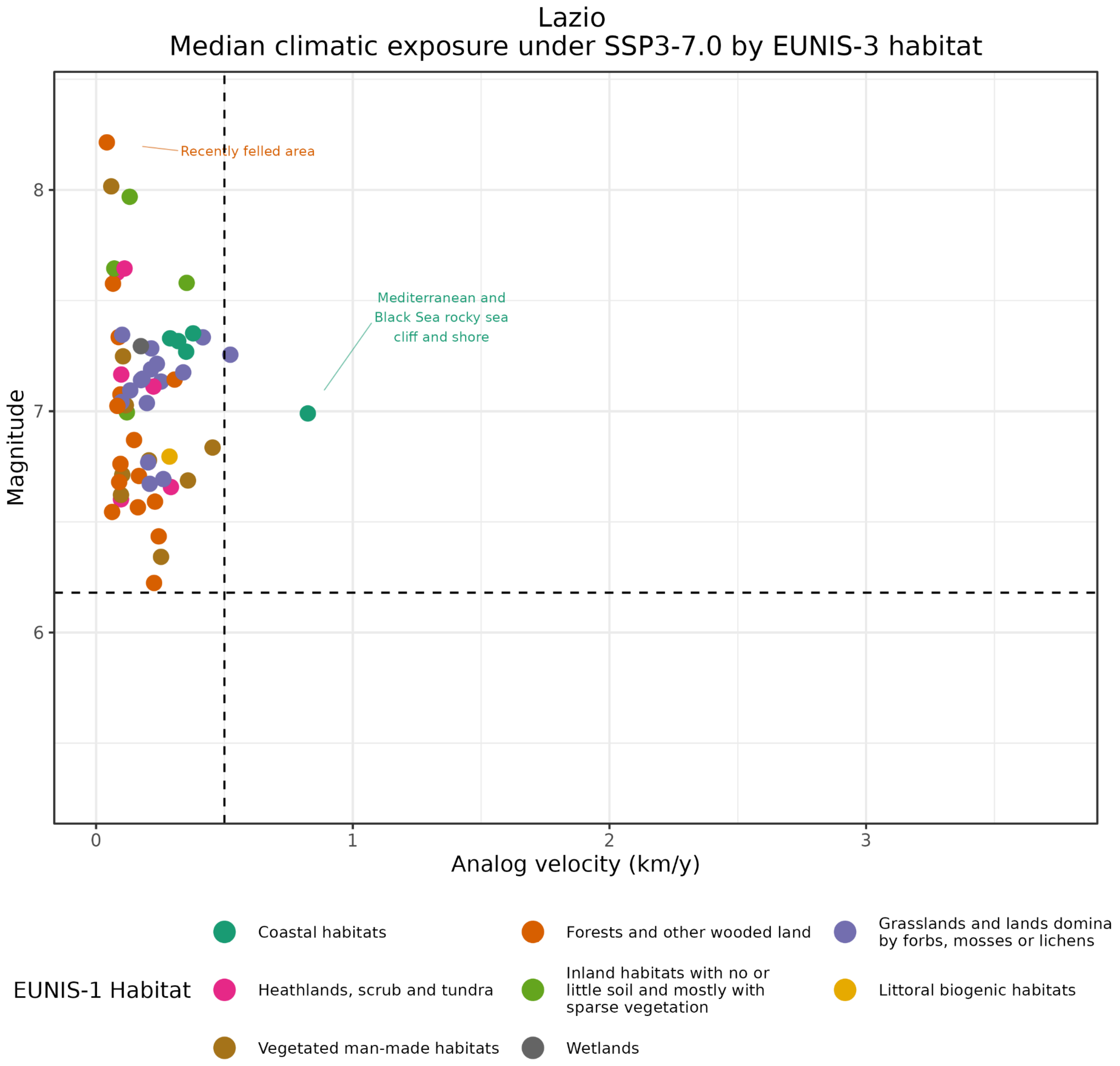

Figure S8. Median climate exposure for each EUNIS level 3 habitat for Lazio under SSP3-7.0

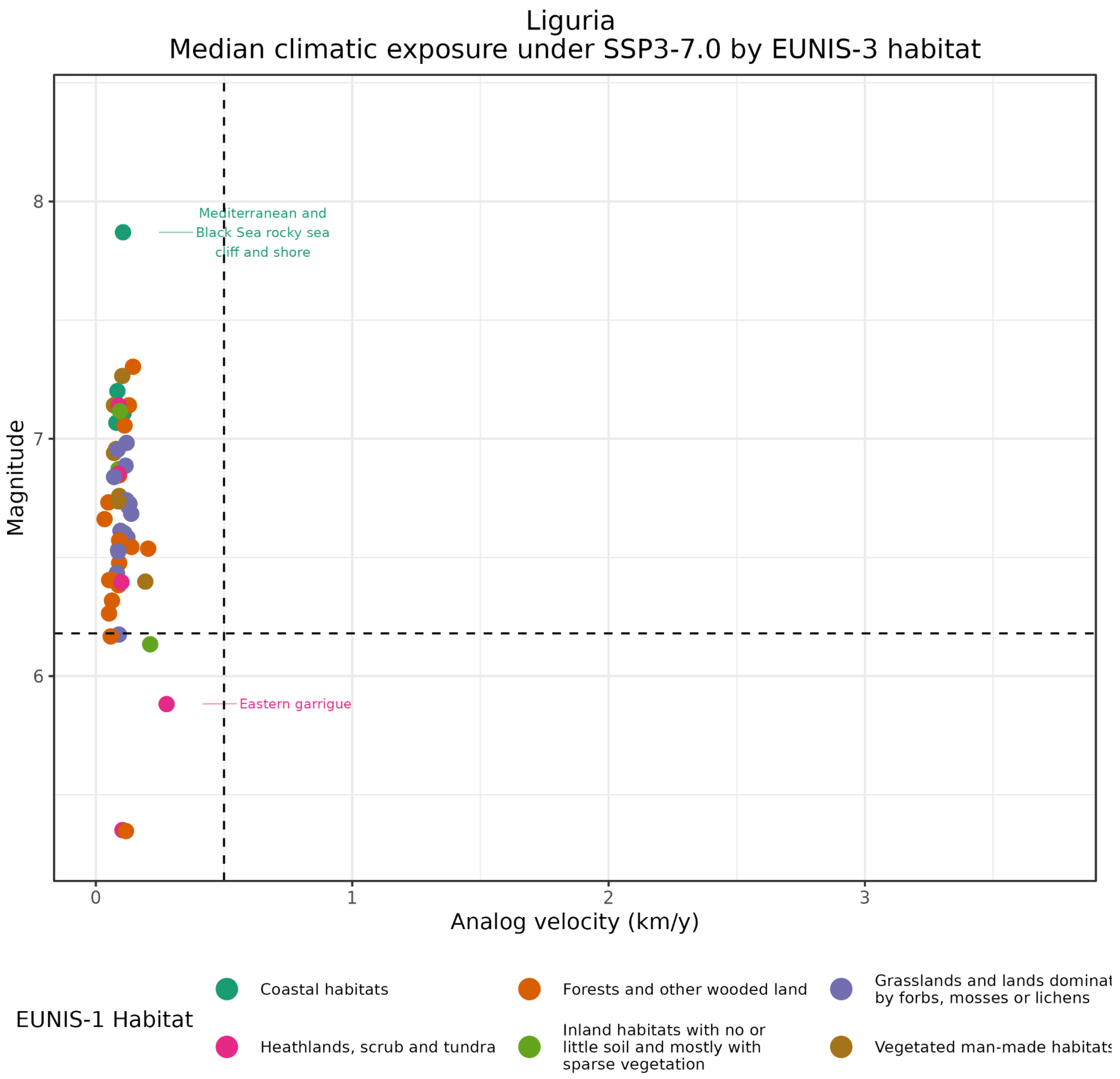

Figure S9. Median climate exposure for each EUNIS level 3 habitat for Liguria under SSP3-7.0

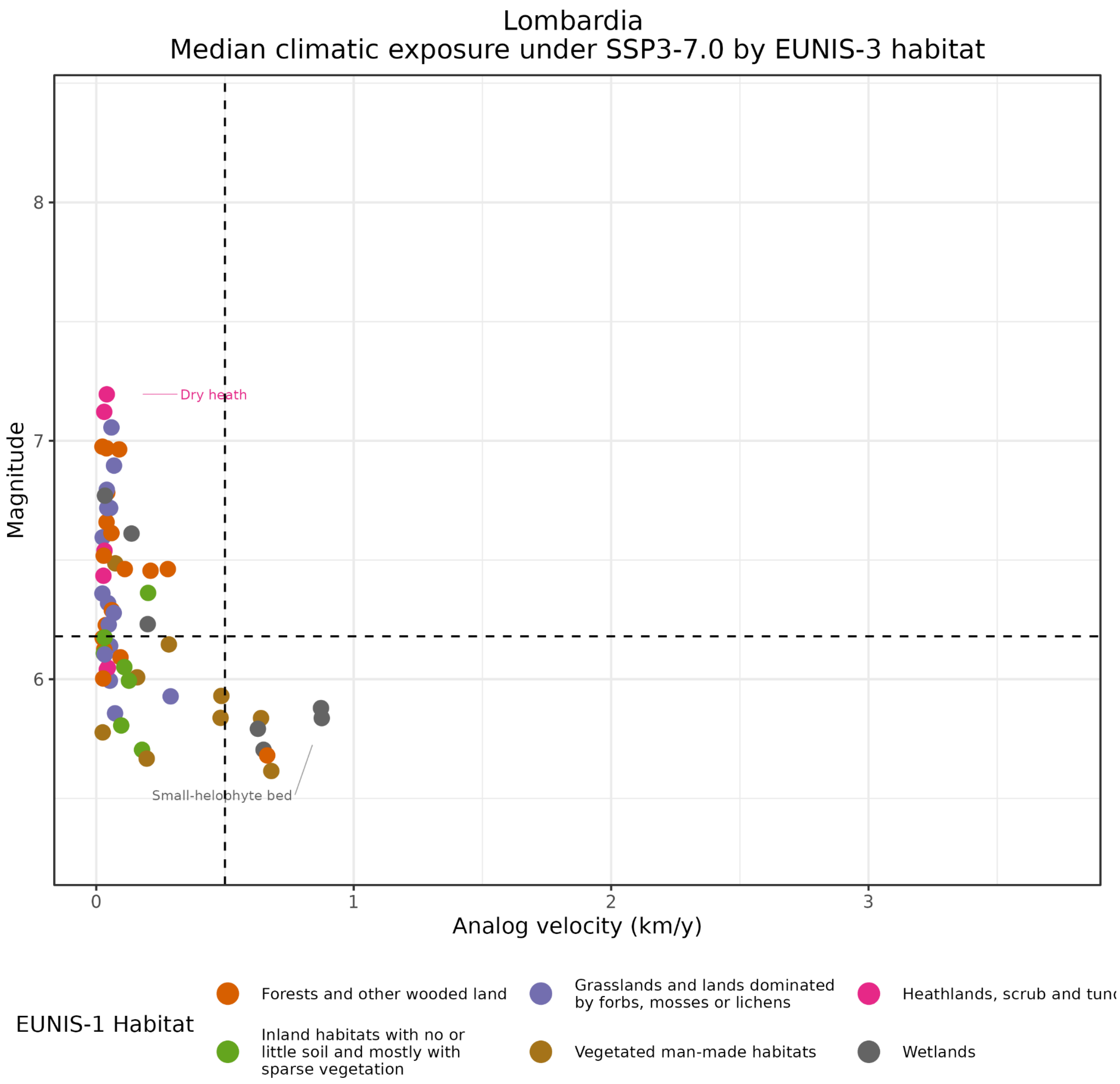

Figure S10. Median climate exposure for each EUNIS level 3 habitat for Lombardia under SSP3-7.0

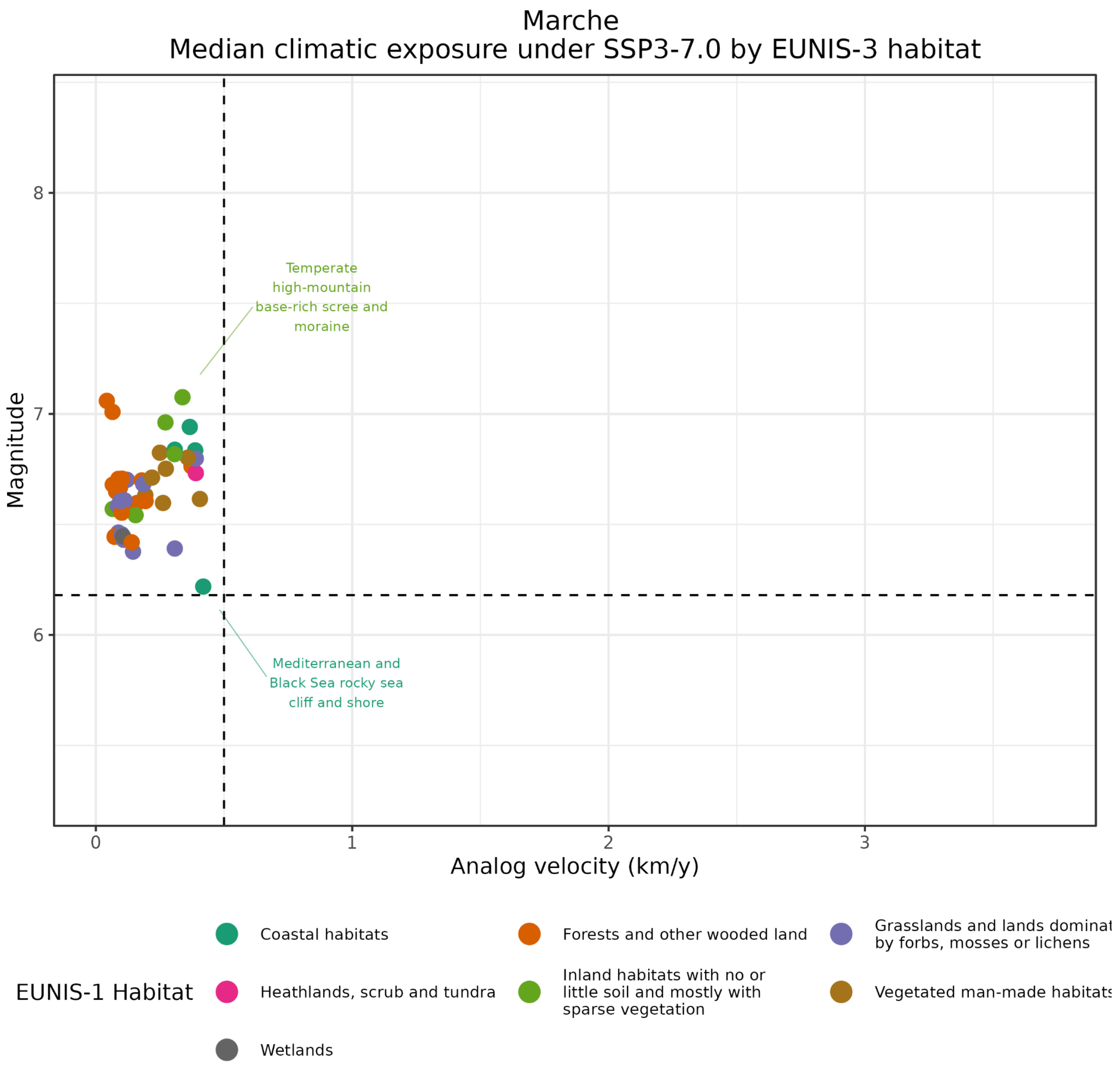

Figure S11. Median climate exposure for each EUNIS level 3 habitat for Marche under SSP3-7.0

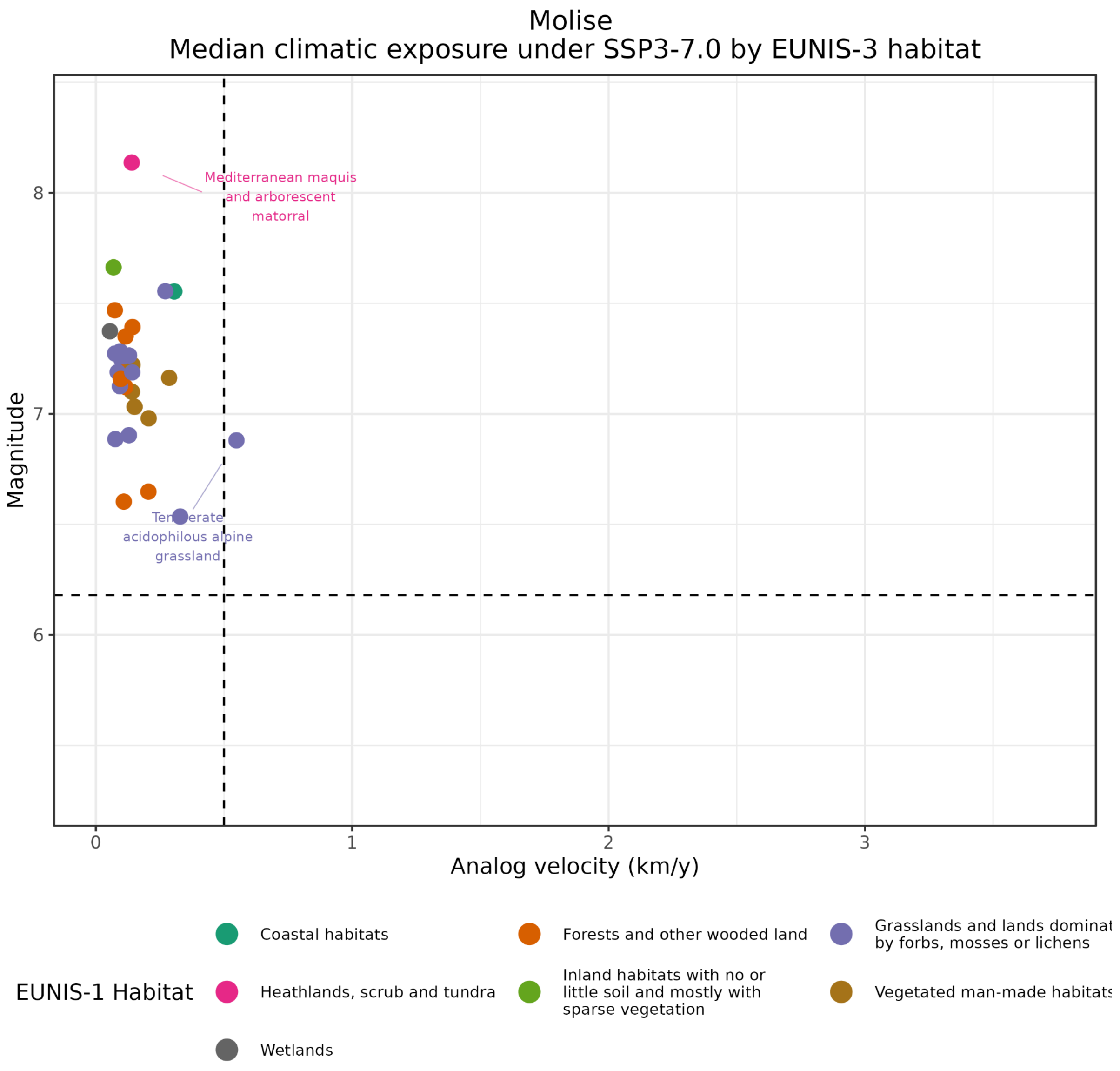

Figure S12. Median climate exposure for each EUNIS level 3 habitat for Molise under SSP3-7.0

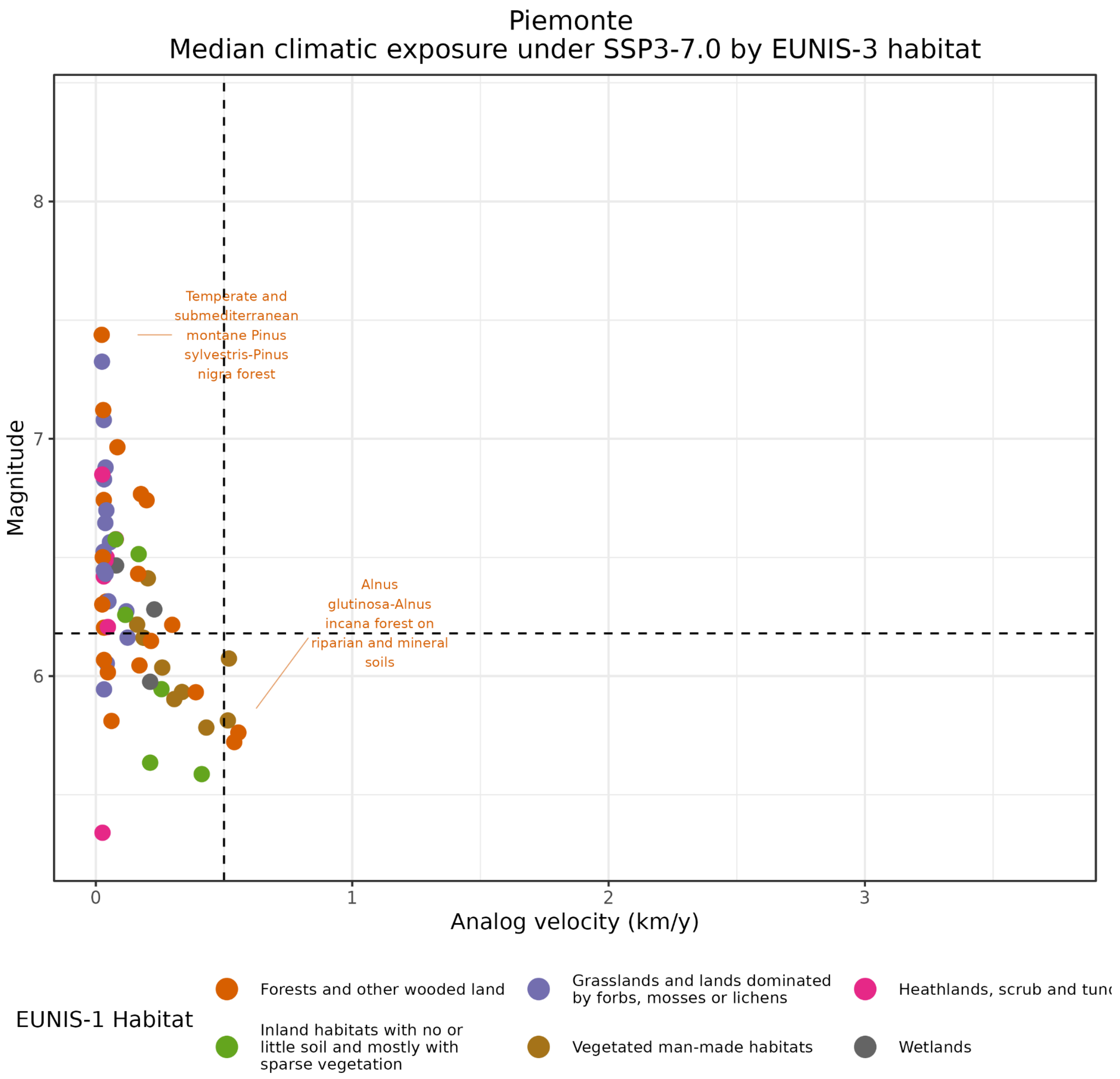

Figure S13. Median climate exposure for each EUNIS level 3 habitat for Piemonte under SSP3-7.0

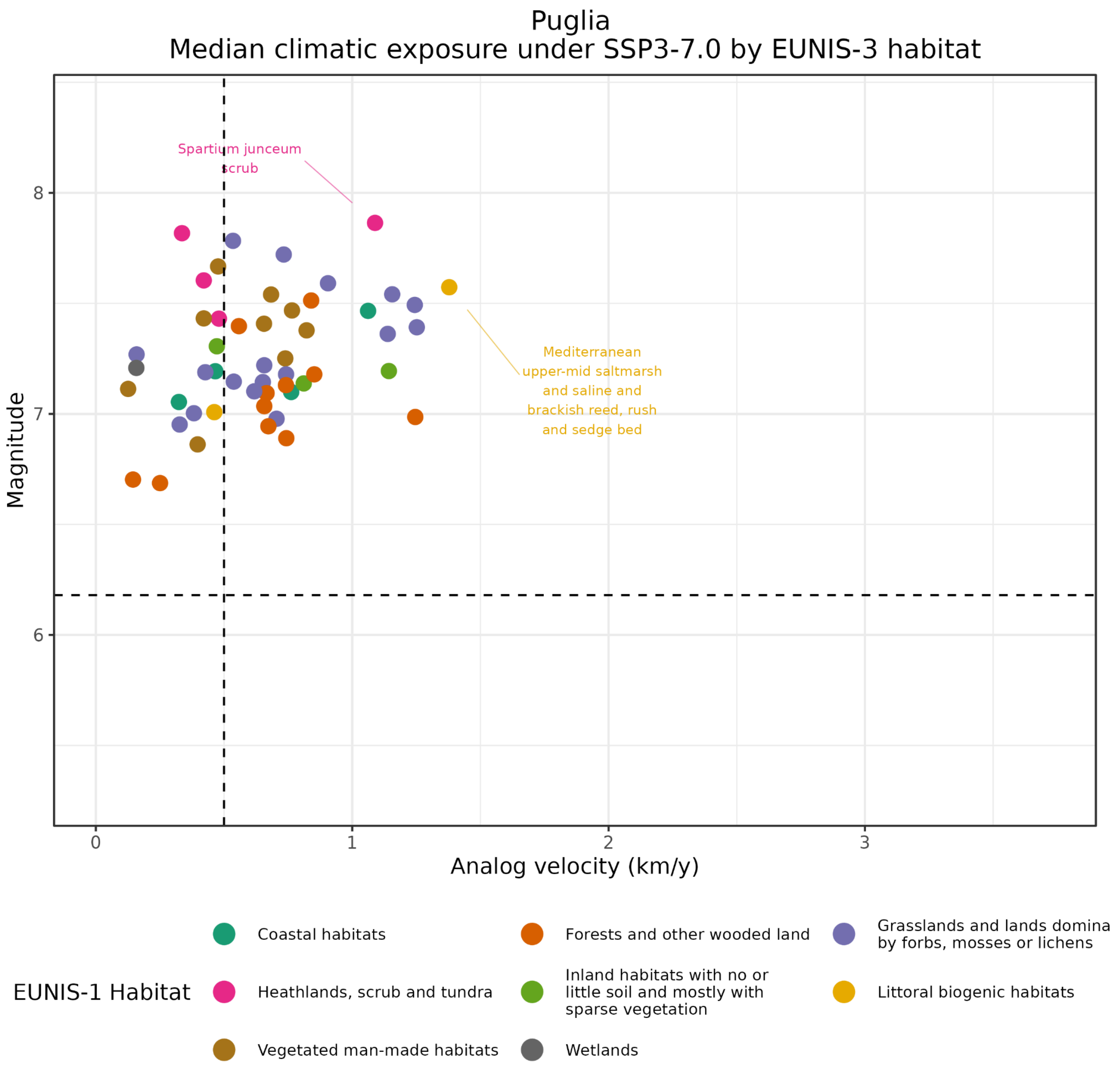

Figure S14. Median climate exposure for each EUNIS level 3 habitat for Puglia under SSP3-7.0

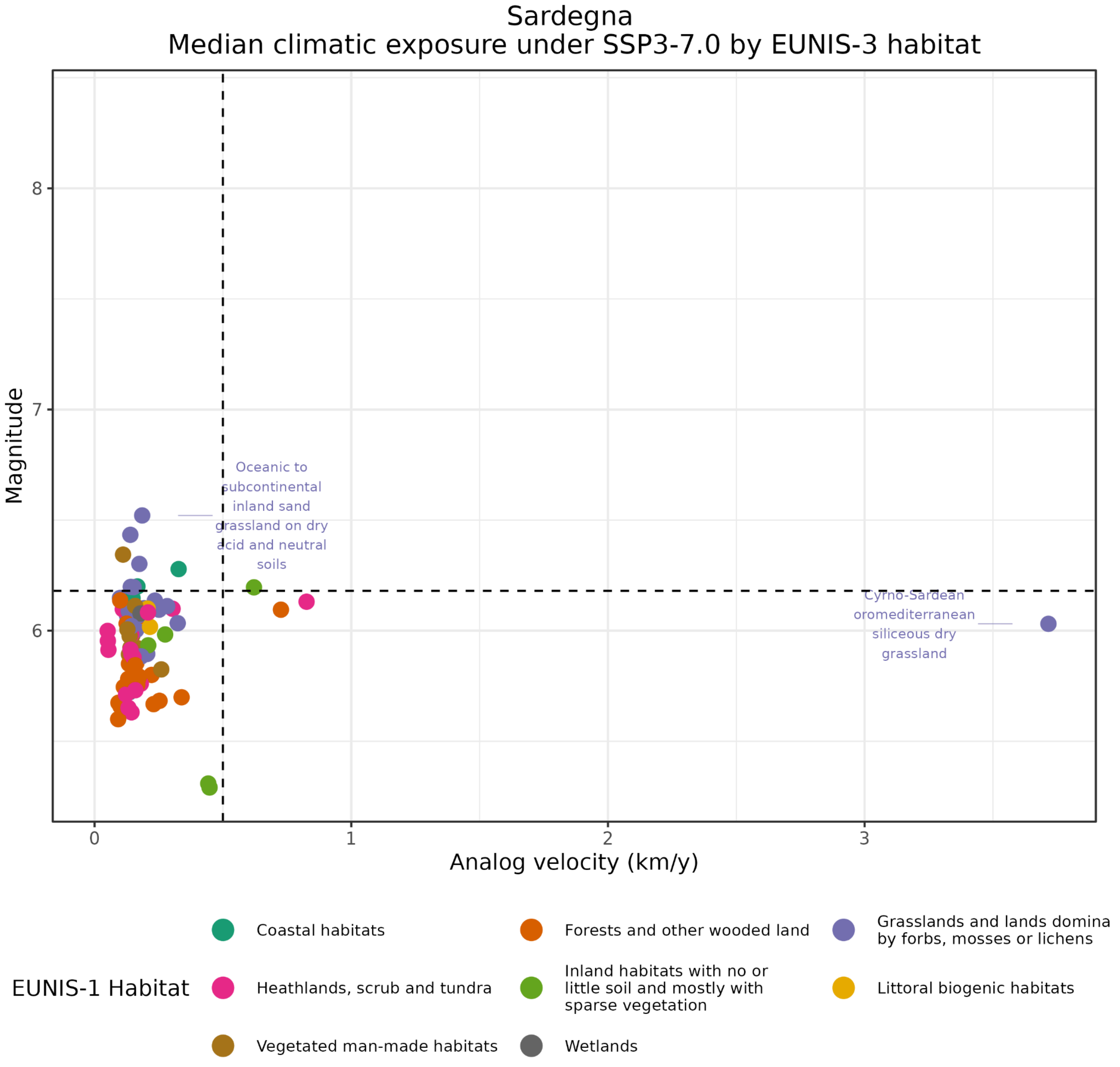

Figure S15. Median climate exposure for each EUNIS level 3 habitat for Sardegna under SSP3-7.0

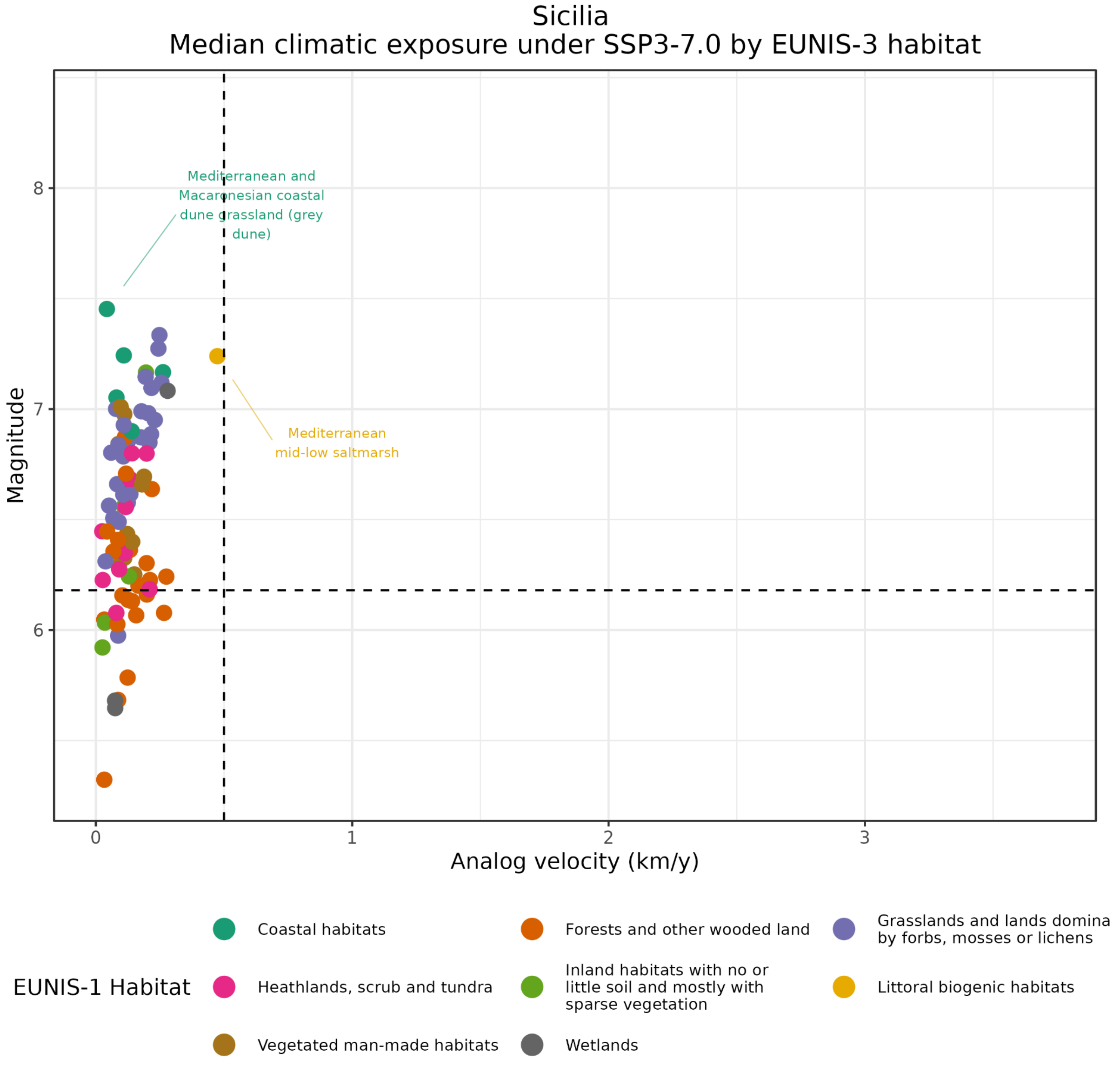

Figure S16. Median climate exposure for each EUNIS level 3 habitat for Sicilia under SSP3-7.0

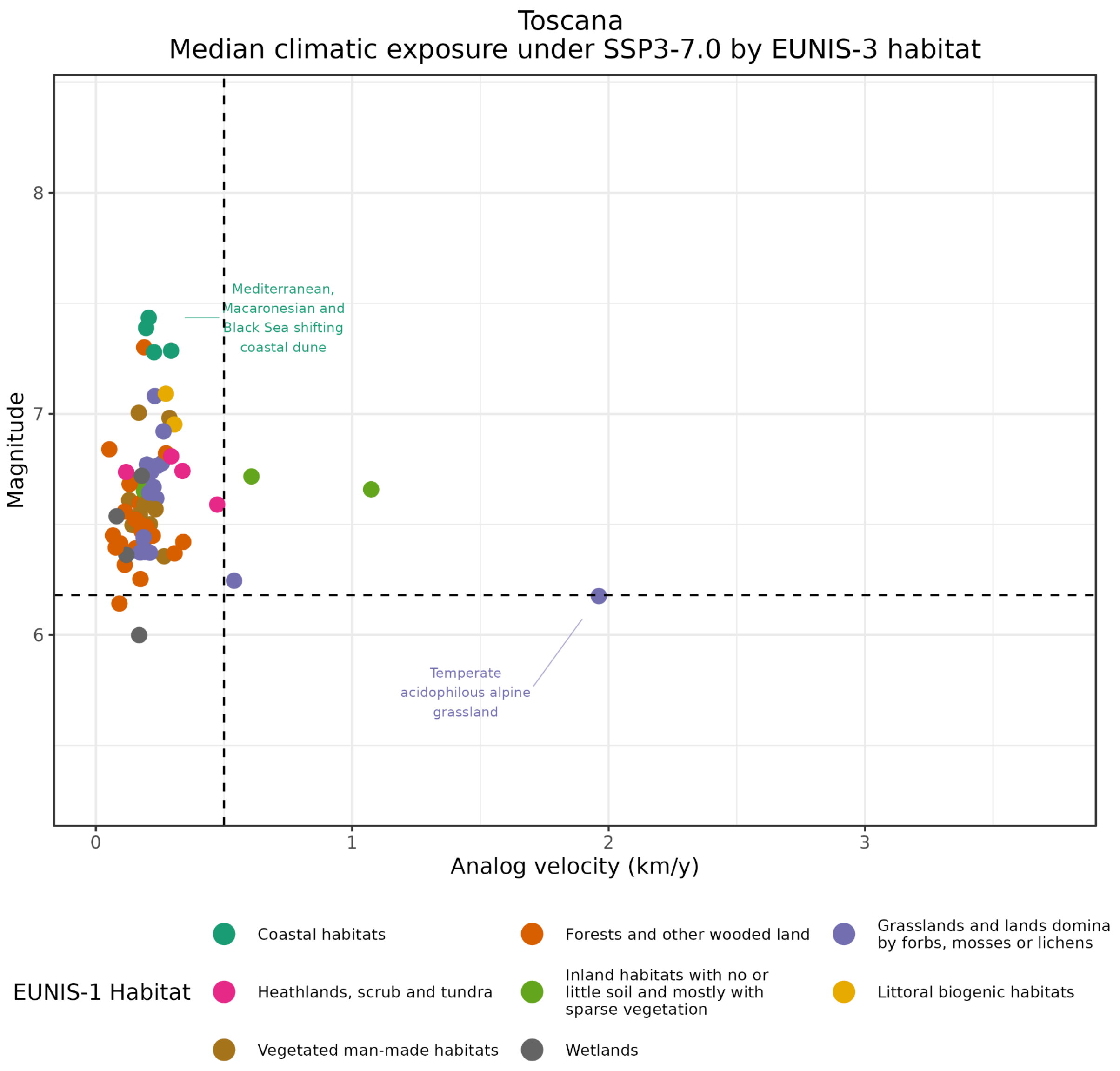

Figure S17. Median climate exposure for each EUNIS level 3 habitat for Toscana under SSP3-7.0

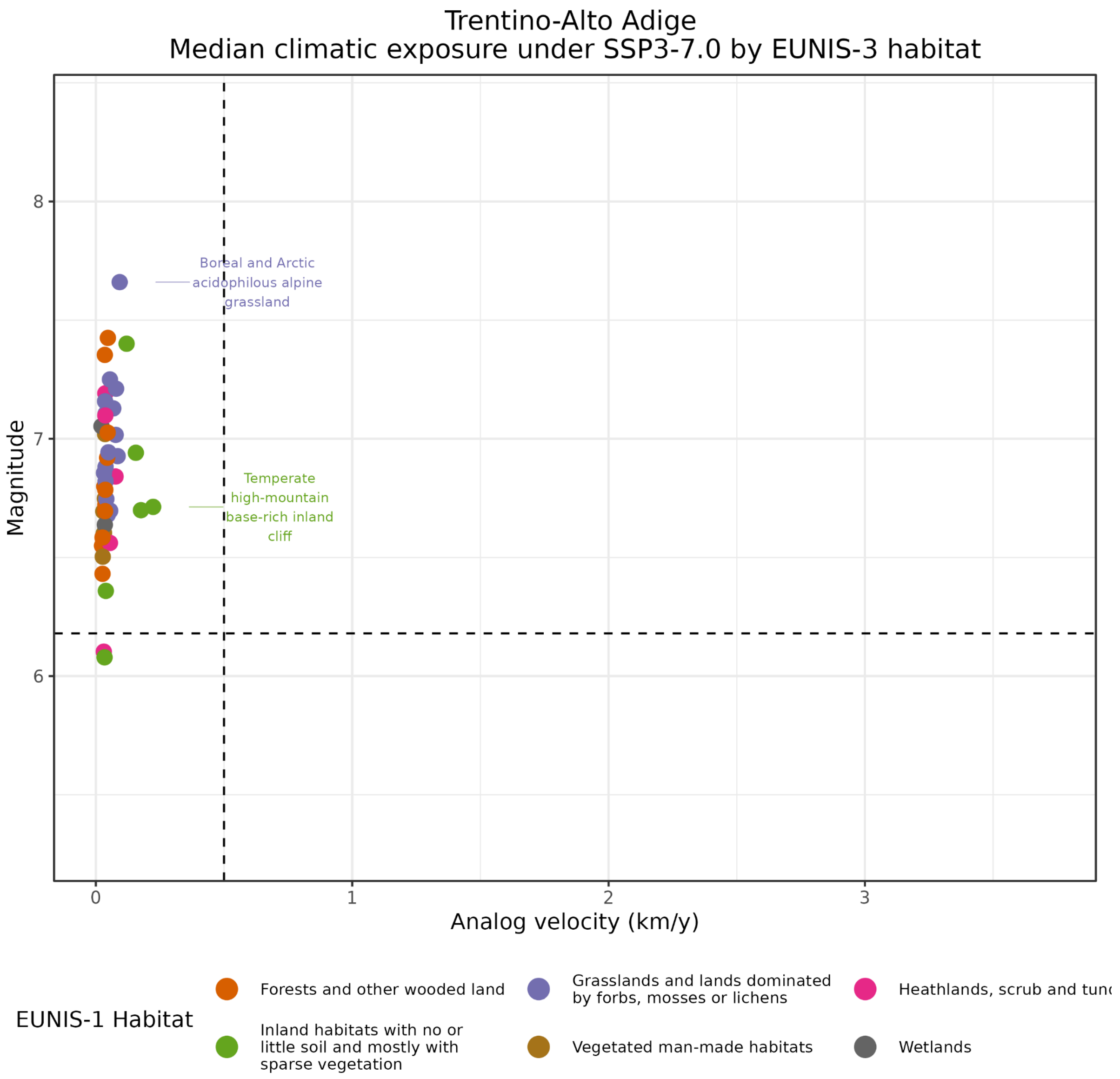

Figure S18. Median climate exposure for each EUNIS level 3 habitat for Trentino-Alto Adige under SSP3-7.0

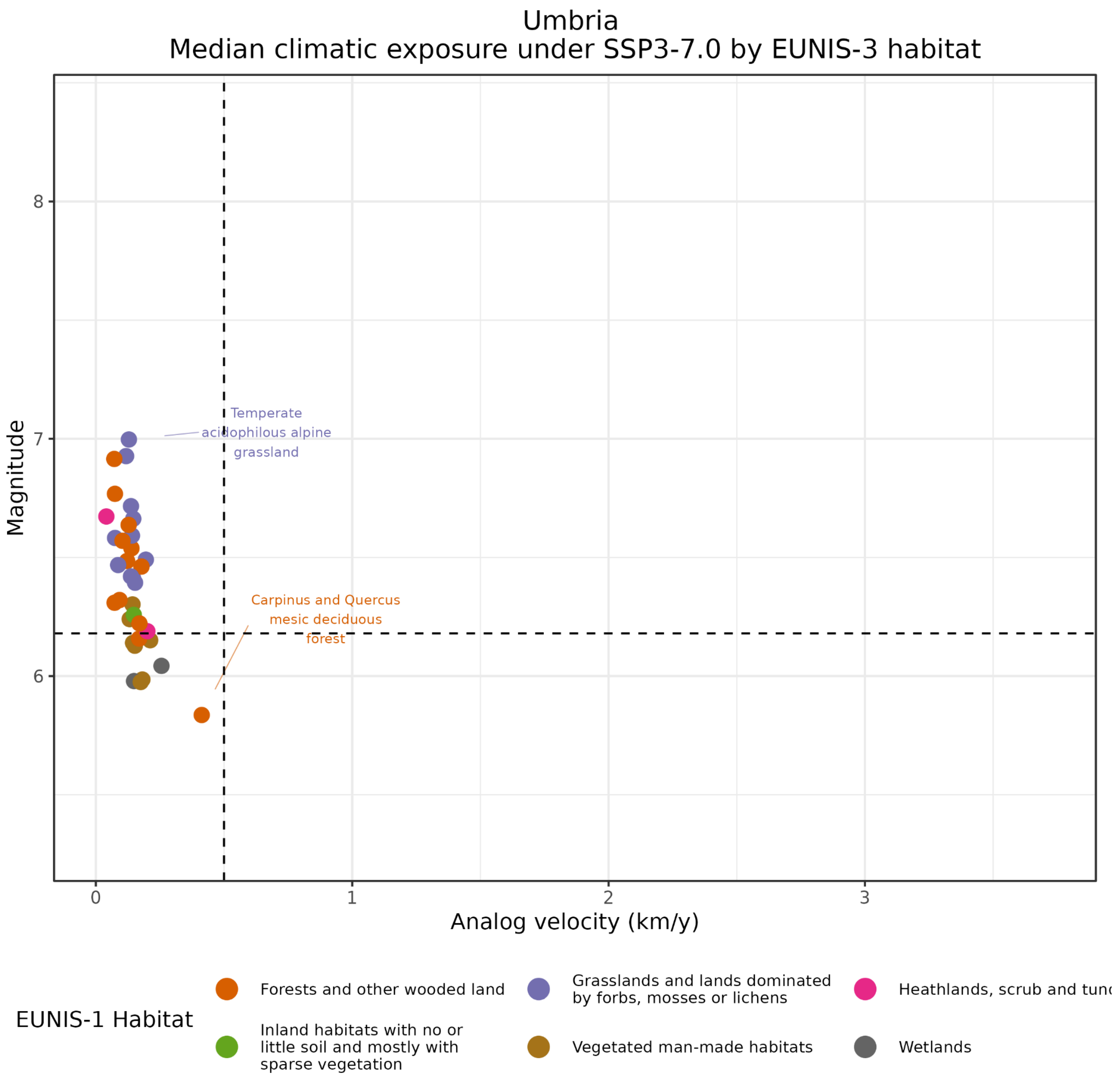

Figure S19. Median climate exposure for each EUNIS level 3 habitat for Umbria under SSP3-7.0

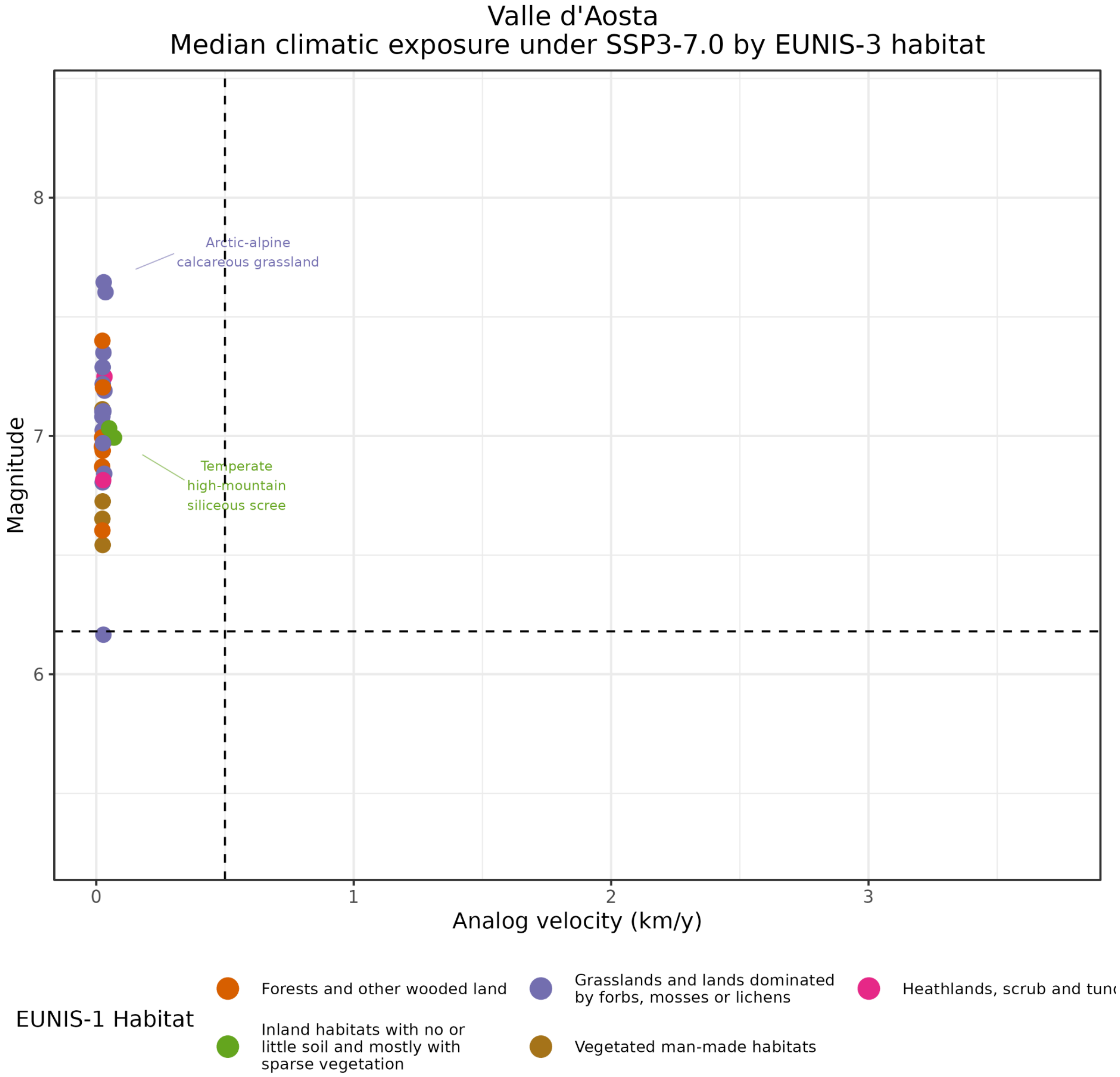

Figure S20. Median climate exposure for each EUNIS level 3 habitat for Valle d'Aosta under SSP3-7.0

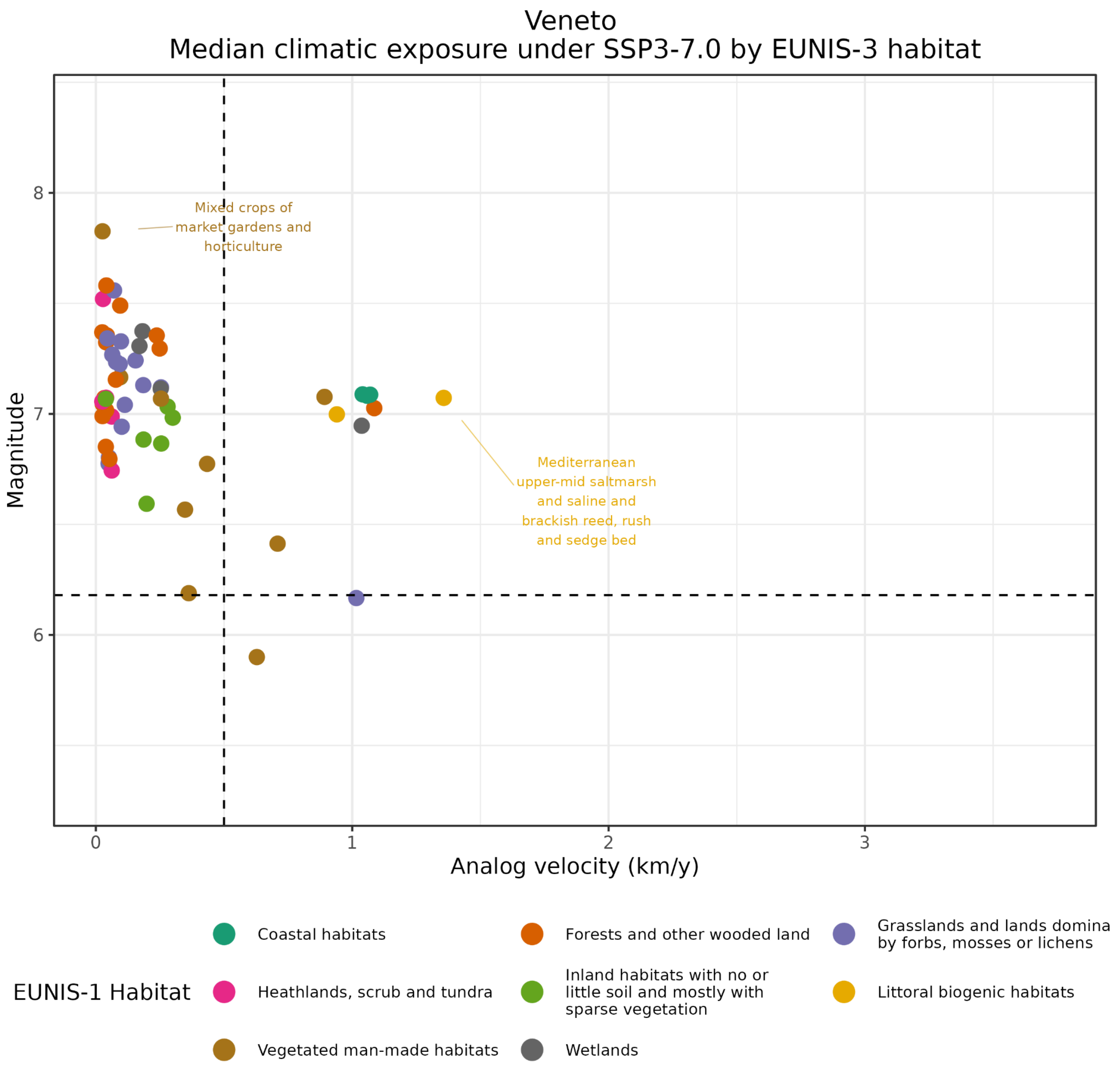


Figure S21. Median climate exposure for each EUNIS level 3 habitat for Veneto under SSP3-7.0
